# Multi ‘omics comparison reveals metabolome biochemistry, not microbiome composition or gene expression, corresponds to elevated biogeochemical function in the hyporheic zone

**DOI:** 10.1101/291096

**Authors:** Emily B. Graham, Alex R. Crump, David W. Kennedy, Evan Arntzen, Sarah Fansler, Samuel O. Purvine, Carrie D. Nicora, William Nelson, Malak M. Tfaily, James C. Stegen

## Abstract

Biogeochemical hotspots are pervasive at terrestrial-aquatic interfaces, particularly within groundwater-surface water mixing zones (hyporheic zones), and they are critical to understanding spatiotemporal variation in biogeochemical cycling. Here, we use multi ‘omic comparisons of hotspots to low-activity sediments to gain mechanistic insight into hyporheic zone organic matter processing. We hypothesized that microbiome structure and function, as described by metagenomics and metaproteomics, would distinguish hotspots from low-activity sediments through a shift towards carbohydrate-utilizing metabolic pathways and elucidate discrete mechanisms governing organic matter processing in each location. We also expected these differences to be reflected in the metabolome, whereby hotspot carbon (C) pools and metabolite transformations therein would be enriched in sugar-associated compounds. In contrast to expectations, we found pronounced phenotypic plasticity in the hyporheic zone microbiome that was denoted by similar microbiome structure, functional potential, and expression across sediments with dissimilar metabolic rates. Instead, diverse nitrogenous metabolites and biochemical transformations characterized hotspots. Metabolomes also corresponded more strongly to aerobic metabolism than bulk C content only (explaining 67% vs. 42% of variation), and bulk C did not improve statistical models based on metabolome composition alone. These results point to organic nitrogen as a significant regulatory factor influencing hyporheic zone organic matter processing. Based on our findings, we propose incorporating knowledge of metabolic pathways associated with different chemical fractions of C pools into ecosystem models will enhance prediction accuracy.

## 1. Introduction

Soils and nearshore sediments contain a vast reservoir of stored carbon. Uncertainty in the fate of these stores is central to constraining future atmospheric CO_2_ concentrations (Burd et al., 2016; Luo et al., 2016; Todd-Brown et al., 2013). Advances in molecular technology has given researchers new ability to characterize mechanisms governing carbon (C) bioavailability, and ultimately the conversion of belowground C pools to CO_2_. These advances are central to ongoing efforts to improve process-based ecosystem models by incorporating microbial and biochemical complexity in biogeochemical processes (Buchkowski et al., 2017; Luo et al., 2016; Wieder et al., 2013; Wieder et al., 2017). Still, there is little consensus on the roles of diverse environmental and microbial factors in enhancing belowground CO_2_ flux predictions (Graham et al., 2016b; Graham et al., 2014; Luo et al., 2016; Rocca et al., 2015; Wieder et al., 2017), and understanding the mechanisms regulating biogeochemistry in natural environments is paramount to conceptualizing the structure and parameterization of models (Luo et al., 2016; Todd-Brown et al., 2013; Wieder et al., 2013).

Recent research has vastly improved both molecular methodologies (Aebersold and Mann, 2003; Gabor et al., 2014; Tfaily et al., 2017; Tringe and Rubin, 2005; Viant, 2008; Wang et al., 2009) and biogeochemical mechanisms represented in ecosystem models (Allison et al., 2010; Wieder et al., 2015a; Wieder et al., 2013; Xu et al., 2014). For instance, parallel advances in environmental chemistry and microbiology now allow for detailed characterization of belowground C pools (e.g., fluorescent methods (Gabor et al., 2014), high-resolution mass spectroscopy (Tfaily et al., 2017), and nuclear magnetic resonance (NMR) (Markley et al., 2017)) as well as the structure and function of environmental microbiomes (Aebersold and Mann, 2003; Tringe and Rubin, 2005; Wang et al., 2009). Newly improved process models account for chemical attributes of C (Liski et al., 2005; Wang et al., 2017), microbial biomass and physiology (Wieder et al., 2014; Wieder et al., 2015a), and nutrient limitations (Wieder et al., 2015b) in their representations of biogeochemistry, among other factors (Verheijen et al., 2015; Wang et al., 2017). Despite these advances, cross-system experimental evidence has revealed many inconsistencies in the importance of different biogeochemical and microbial properties explaining respiration rates (Graham et al., 2016b; Graham et al., 2017b; Hartman et al., 2017), suggesting a need for greater spatially-explicit understanding of mechanisms involved in organic matter processing (Graham et al., 2017b; Krause et al., 2017).

Terrestrial-aquatic interfaces are recognized active regions of C metabolism, and areas of subsurface groundwater-surface water mixing (hyporheic zones) in particular are critical in determining the fate of organic matter globally (Battin et al., 2009; Boulton et al., 1998; Cole et al., 2007; Marín-Spiotta et al., 2014; Regnier et al., 2013). Hyporheic zones contribute disproportionately to stream and river metabolism (Gomez-Velez et al., 2015; Huizenga et al., 2017; Stegen et al., 2016; Stern et al., 2017), with up to 96% of ecosystem metabolism occurring in this zone in some headwater systems (Naegeli and Uehlinger, 1997). Hotspots and hot moments of enhanced biogeochemical activity at confined locations or time points are common within the hyporheic zone (Boulton et al., 1998; Krause et al., 2017; McClain et al., 2003). These hotspots often correspond to vegetation patterns (Harms and Grimm, 2008; McClain et al., 2003; Schade et al., 2001) and understanding their drivers is critical to accurately representing organic matter decomposition within ecosystem models.

Our objective is to determine molecular mechanisms associated with hotspots of aerobic metabolism in the hyporheic zone. Previous work in this system has shown differences in (1) aerobic metabolism across variable mixing of water bodies with distinct C chemistries (Stegen et al., 2016; Stegen et al., 2018) and (2) metabolic processes associated with C oxidation in the presence or absence of vegetation (Graham et al., 2017b). Recent work has also shown that elevated rates of metabolism globally correspond to carbohydrate metabolism in soil metagenomes (Hartman et al., 2017). Based on this research, we hypothesize that microbiome structure and function support elevated carbohydrate metabolism in hyporheic zone hotspots, resulting in higher rates of aerobic metabolism. We expect this dynamic to be reflected in sediment metagenomes, metaproteomes, and metabolomes, whereby hotspot microbiome composition, protein expression, and metabolites are enriched in sugar-associated pathways and compounds, relative to low-activity sediments.

## 2. Materials and Methods

### 2.1. Site Description

This study was conducted along the Columbia River shoreline within the Hanford 300 Area (approximately 46° 22’ 15.80”N, 119° 16’ 31.52”W) in eastern Washington State (Graham et al., 2016a; Graham et al., 2017a; Graham et al., 2017b; Slater et al., 2010; Zachara et al., 2013). The Columbia River experiences shoreline geographic variation in vegetation patterns, substrate geochemistry, microbiome composition, and rates of biogeochemical activity (Arntzen et al., 2006; Graham et al., 2017b; Lin et al., 2012; Peterson and Connelly, 2004; Slater et al., 2010; Stegen et al., 2016; Stegen et al., 2012; Zachara et al., 2013); constituting an ideal system for examining mechanisms associated with biogeochemical hotspots.

Liquid N_2_-frozen sediment profiles (0-60 cm) were collected along two shoreline transects perpendicular to the Columbia River in March 2015, separated by a distance of ∼170m. We collected profiles at three locations in each transect with 5m spacing within a spatial domain of ∼175 x 10m. In each transect, the lower bank profile was located at ∼0.5m (vertical distance) below the water line and the upper bank profile was located ∼0.5m (vertical distance) above the water line (approximately 10m horizontal distance), with the third profile situated at the midpoint. Each profile was sectioned into 10-cm intervals from 0-60cm. To capture a range of biogeochemical activities, one transect had dense vegetation, consisting of a closed canopy of woody perennials *Morus rubra* (Red Mulberry) and *Ulmus rubra* (Slippery Elm), and one transect was characterized by a cobbled armor layer with virtually no vegetation.

### 2.2. Sample Collection

Liquid N_2_-frozen sediment profiles were collected as outlined in Moser et al. (2003) using a method modified from Lotspeich and Reed (1980) and Rood and Church (1994). A pointed stainless steel tube (152 cm length, 3.3 cm outside diameter, 2.4 cm inside diameter) was driven into the river bed to a depth of ∼60cm. Liquid N_2_ was poured down the tube for ∼15 minutes, until a sufficient quantity of material had frozen to the outside of the tube. The tube and attached material were removed from the riverbed with a chain hoist suspended beneath a tripod. Profiles were placed over an aluminum foil lined cooler containing dry ice. Frozen material was removed with a mallet. The material was then wrapped in the foil and transported on dry ice to storage at −80°C. In the lab, profiles were sectioned into 10cm depth intervals from 0-60 cm (n = 6 per profile, except for the unvegetated upper bank profile which was sectioned only from 30-60cm; total n = 33)

### 2.3. Physicochemistry

Details concerning physicochemical assays are provided in the Supporting Information. Briefly, we determined the particle distribution of sediments by separating size fractions via sieving; total nitrogen, sulfur, and carbon content were determined using an Elementar vario EL cube (Elementar Co.Germany); NH_4_^+^ was extracted with KCl and measured with Hach Kit 2604545 (Hach, Loveland, Co); iron content was measured with a ferrozine assay; and all other ion concentrations were measured by inductively coupled plasma mass spectrometry (ICP-MS) on HCl extractions. Aerobic metabolism was determined with a resazurin reduction assay, modified from *Haggerty et al.* (2009). Data are provided in Fig. S1-S3.

### 2.4. FT-ICR-MS solvent extraction and data acquisition

We leverage state of science chemical extraction protocols combined with Electrospray ionization (ESI) and Fourier transform ion cyclotron resonance (FT-ICR) mass spectrometry (MS) to infer differences in metabolites among our samples. ESI FT-ICR-MS introduces intact organic molecules into the MS without fragmentation and allows for the detection of a wide range of chemical compounds (Tfaily et al., 2015; Tfaily et al., 2017). The use of 12 Tesla (T) FT-ICR-MS offers high mass resolving power (>1M) and mass measurement accuracy (<1 ppm), and while nascent in its application within complex environmental systems, it has emerged as a robust method for determining the chemistry of natural organic compounds (Kim et al., 2003; Koch et al., 2005; Tfaily et al., 2011; Tremblay et al., 2007). Additionally, Tfaily et al. (2015; 2017) have optimized metabolite characterization from soils and sediments by sequential extraction with polar and non-polar solvents tailored to the sample set of interest. Tfaily’s extraction procedures have been coupled to ESI FT-ICR-MS to distinguish metabolites among ecosystems and soil types (Tfaily et al., 2015; Tfaily et al., 2017) as well as to provide information on the utilization of distinct metabolites among samples within a single environment (Bailey et al., 2017; Graham et al., 2017b; Stegen et al., 2018).

Here, we used three solvents with different polarities ––water (H_2_O), methanol (CH_3_OH, ‘MeOH’) and chloroform (CHCl_3_)––to sequentially extract a large diversity of organic compounds from samples, according to Tfaily et al. (2015; 2017). Water extractions were performed first, followed by MeOH and then CHCl_3_. Previous work has shown that each solvent is selective towards specific types of compounds (Tfaily et al., 2015) and that combining peaks from all extractions provides a more comprehensive description of metabolite composition than any single extraction procedure independently (Tfaily et al., 2017). Water is a polar solvent with a selection bias for carbohydrates with high O/C ratios, amino-sugars, and other labile polar compounds (Tfaily et al., 2015). Conversely, CHCl_3_ is selective for non-polar lipids associated with mineral interactions and cellular membranes (Tfaily et al., 2015). Methanol has a polarity in between that of water and CHCl_3_ and extracts a mixture of compounds that water and CHCl_3_ extract (Tfaily et al., 2015). Ultra-high resolution mass spectrometry of the three different extracts from each sample was carried out using a 12 Tesla Bruker SolariX FT-ICR-MS located at the Environmental Molecular Sciences Laboratory (EMSL) in Richland, WA, USA. An expanded description of extraction procedures; instrument calibration and maintenance; and sample injection is presented in the Supplemental Material.

### 2.5. FT-ICR-MS data processing

One hundred forty-four individual scans were averaged for each sample and internally calibrated using an organic matter homologous series separated by 14 Da (–CH_2_ groups). The mass measurement accuracy was less than 1 ppm for singly charged ions across a broad *m*/*z* range (100-1200 *m*/*z*). The mass resolution was ∼ 350K at 339 m/z. Data Analysis software (BrukerDaltonik version 4.2) was used to convert raw spectra to a list of m/z values applying FTMS peak picker module with a signal-to-noise ratio (S/N) threshold set to 7 and absolute intensity threshold to the default value of 100.

For each sample, we combined peaks detected in all three extractions to yield metabolite composition. Peak data were treated as presents/absence data, where a peak was considered to be present in a sample if it was present in at least one of H_2_O-, MeOH-, or CHCl_3_-extracted samples. Presence/absence data were used because peak intensity differences are reflective of ionization efficiency as well as relative abundance (Kujawinski and Behn, 2006; Minor et al., 2012; Tfaily et al., 2015; Tfaily et al., 2017).

Putative chemical formulae were then assigned using in-house software following the Compound Identification Algorithm (CIA), proposed by Kujawinski and Behn (2006), modified by Minor et al. (2012), and previously described in Tfaily et al. (2017). Chemical formulae were assigned based on the following criteria: S/N >7, and mass measurement error <1 ppm, taking into consideration the presence of C, H, O, N, S and P and excluding other elements. To ensure consistent formula assignment, we aligned all sample peak lists for the entire dataset to each other in order to facilitate consistent peak assignments and eliminate possible mass shifts that would impact formula assignment. We implemented the following rules to further ensure consistent formula assignment: (1) we consistently picked the formula with the lowest error and with the lowest number of heteroatoms and (2) the assignment of one phosphorus atom requires the presence of at least four oxygen atoms.

The chemical character of thousands of peaks in each sample’s ESI FT-ICR-MS spectrum was evaluated on van Krevelen diagrams. Compounds were plotted on the van Krevelen diagram on the basis of their molar H:C ratios (y-axis) and molar O:C ratios (x-axis) (Kim et al., 2003). Van Krevelen diagrams provide a means to visualize and compare the average properties of organic compounds and assign compounds to the major biochemical classes (e.g., lipid-, protein-, lignin-, carbohydrate-, and condensed aromatic-like).

### 2.6. Identification of putative biochemical transformations using FT-ICR-MS

To identify potential biochemical transformations, we followed the procedure detailed by Breitling *et al*. (2006) and employed by Bailey *et al*. (2017), Graham *et al*. (2017b), and Stegen *et al*. (2018). The mass difference between m/z peaks extracted from each spectrum with S/N>7 were compared to commonly observed mass differences associated with biochemical transformations. All possible pairwise mass differences were calculated within each extraction type for each sample, and differences (within 1ppm) were matched to a list of 92 common biochemical transformations (e.g., gain or loss of amino groups or sugars, Table S1). For example, a mass difference of 99.07 corresponds to a gain or loss of the amino acid valine, while a difference of 179.06 corresponds to the gain or loss of a glucose molecule. Pairs of peaks with a mass difference within 1 ppm of our transformation list were considered to be related by the corresponding compound. This approach is feasible with FT-ICR-MS data because the set of peaks in each sample are related by measureable and clearly defined mass differences corresponding to gains and losses of compounds.

### 2.7. Metagenome sequencing and annotation

To release biomass from sediment particles, thawed samples were suspended in 20mL of chilled PBS/0.1% Na-pyrophosphate solution and vortexed for 1 min. The suspended fraction was decanted to a fresh tube and centrifuged for 15’ at 7000 x *g* at 10°C. DNA was extracted from the resulting pellets using the MoBio PowerSoil kit in plate format (MoBio Laboratories, Inc., Carlsbad, CA) following manufacturer’s instructions, with the addition of a 2-hour proteinase-K incubation at 55°C prior to bead-beating to facilitate cell lysis.

Purified genomic DNA was submitted to the Joint Genome Institute under JGI/EMSL proposal 1781 for paired-end sequencing on an Illumina HiSeq 2500 sequencer (Table S2). Reads were processed by BBDuk to remove adapters (ktrim=r, minlen=40, minlenfraction=0.6, mink=11, tbo, tpe, k=23, hdist=1, hdist2=1, ftm=5) and trim for quality <12 (maq=8, maxns=1, minlen=40, minlenfraction=0.6, k=27, hdist=1, trimq=12, qtrim=rl) and screened for contaminants against a masked version of human HG19 using BBMap (fast local minratio=0.84 maxindel=6 tipsearch=4 bw=18 bwr=0.18 usemodulo printunmappedcount idtag minhits=1). (BBDuk and BBMap are available at https://sourceforge.net/projects/bbmap) Remaining reads were assembled with megahit (Li et al., 2016) using default parameters and k-list=23,43,63, 83,103,123. Gene calling and functional and taxonomic annotation was performed by MGAP v4.11.4 (Huntemann et al., 2015). Data sets are available through the JGI Genome Portal (http://genome.jgi.doe.gov). Project identifiers are listed in Table S2.

### 2.8. Metaproteomics

Sediment samples were prepared for proteome analysis as per Nicora et al. (in press) and detailed below. Additional details are provided in the Supplemental Material.

Proteins were extracted from 30 g of lyophilized sediment using MPLex direct SDS buffer extraction. Sediment was weighed into 50 mL screw-cap self-standing tubes (Next Advance, Averill Park, NY) along with 0.9-2.0 mm stainless steel beads, 0.1 mm zirconia beads and 0.1 mm garnet beads and 8 mL of 60% MeOH in nanopure water. The samples were all bead beat in a 50mL Bullet Blender (Next Advance, Averill Park, NY) at speed 12 for 15 minutes at 4°C and transferred into chemical compatible polypropylene 50 mL tubes (Olympus Plastics, Waltham, MA). The dirty tube was rinsed with 2 mL of 60% MeOH and combined with the sample along with 12 mL of ice-cold chloroform and probe sonicated at 60% amplitude for 30 seconds on ice, allowed to cool on ice and sonicated again. Samples were incubated for 5 min at-80°C, vortexed for 1 min and centrifuged at 4,500x g for 10 min at 4°C. The upper aqueous phase was collected into a large glass vial, being careful not to touch the protein interphase. The interphase was collected using a large flat spatula into a separate tube and 5mL of ice cold 100% methanol was added to the protein, vortexed and centrifuged at 4,500x g for 5 mins at 4°C to pellet the protein. The supernatant was decanted off and the protein allowed to dry upside down on a Kim-wipe. Meanwhile, the bottom organic phase was collected into a separate large glass vial and 5mL of nanopure water was added to the large soil particulates along with 25mL of cold (−20 °C) chloroform:methanol (2:1, v:v) solution and probe sonicated for 30 seconds as described previously. The sample was allowed to cool at −20°C and centrifuged at 4,500x g for 10 min at 4°C. The upper aqueous phase metabolites and bottom organic phase lipids were collected together with the supernatant metabolites and lipids then dried down completely in a vacuum concentrator (Labconco, Kansas City, MO) and stored at −20 °C for analysis. The dirty protein interphase had 20 mL of an SDS-Tris buffer (4% SDS, 100mM DTT in 100 mM Tris-HCl, pH 8.0) added and probe sonicated at 20% amplitude to bring into solution and incubated at 95°C for 5 minutes to solubilize the protein and allowed to cool for 20 mins at 4°C. The samples were centrifuged at 4,500x g for 10 mins and the supernatants were decanted into chemical compatible polypropylene 50 mL tubes. The proteins were precipitated by adding up to 20% trichloroacetic acid (TCA), vortexed and placed in a −20°C freezer for 1.5 hrs. The samples were thawed and centrifuged at 4,500x g at 4°C for 10 mins to collect the precipitated protein. The supernatant was gently decanted into waste and 2 mL of ice cold acetone was added to the pellet and re-suspend by vortexing. The sample was placed at −80°C for ∼5 mins to ensure the samples were cold and centrifuged for 10 mins at 4,500x g at 4°C. The acetone was removed by gently pouring into waste. The pellets were allowed to dry inverted on a Kim Wipe for ∼15 mins. The protein interphases from the supernatants and the protein interphases from the particulates were combined using 100-200ul of SDS-Tris buffer and FASP digested (described in the Supplemental Material).

Extractions were analyzed on a Q-Exactive Plus mass spectrometer (Thermo Electron, Waltham, MA) coupled to a Waters NanoAcquity high performance liquid chromatography systems (Waters Corporation, Milford, MA) through 75 um x 70 cm columns packed with Phenomenex Jupiter C-18 derivatized 3 um silica beads (Phenomenex, Torrance, CA). Samples were loaded onto columns with 0.05% formic acid in water and eluted with 0.05% formic acid in Acetonitrile over 100 minutes. Ten data-dependent MS/MS scans (17.5K resolution, centroided) were recorded for each survey MS scan (35K resolution) using normalized collision energy of 30, isolation width of 2.00, and rolling exclusion window of +/-1 Th lasting 30 seconds before previously fragmented signals are eligible for re-analysis.

For protein identification, a single reference protein file was developed from the combination of 33 metagenome files (‘assembled.faa’ and ‘product_names’ files) available from the Joint Genome Institute (JGI Project IDs and related download links in Supplemental Material). Exact sequence duplicates were removed, and 16 commonly observed contaminants (e.g. tryptic fragments, human keratins, and serum albumin precursors) were included. Final file contained 10,063,272 protein entries from 1,299,102,456 amino acids, 2.04GB in size.

The MS/MS spectra from all LC-MS/MS datasets were converted to ASCII text (.dta format) using MSConvert (http://proteowizard.sourceforge.net/tools/msconvert.html) which more precisely assigns the charge and parent mass values to an MS/MS spectrum. The data files were then interrogated via target-decoy approach (http://www.ncbi.nlm.nih.gov/pubmed/20013364) using MSGFPlus (http://www.ncbi.nlm.nih.gov/pubmed/25358478) using a +/-20 ppm parent mass tolerance, partially tryptic digestion enzyme settings, and a variable posttranslational modification of oxidized Methionine. All MS/MS search results for each dataset were collated into tab separated ASCII text files listing the best scoring identification for each spectrum.

Collated search results were further combined into a single result file. These results were imported into a Microsoft SQL Server database. Results were filtered to >1% FDR using an MSGF+ supplied Q-Value that assesses reversed sequence decoy identifications for a given MSGF score across each dataset. Using the protein references as a grouping term, unique peptides belonging to each protein were counted, as were all peptide spectrum matches (PSMs) belonging to all peptides for that protein (i.e. a protein level observation count value). PSM observation counts reported for each sample that was analyzed. Cross-tabulation tables were created to enumerate protein level PSM observations for each sample, allowing low-precision quantitative comparisons to be made. Identified proteins were searched against dbCAN v5 (Yin et al., 2012) using hmmer v3.1 (Finn et al., 2011). CAZy families were assigned based on cutoff suggestions of the maintainers of dbCAN. We removed singletons and proteins not present in at least 25% of samples prior to statistical analysis. Samples with less than 10 total protein counts (n = 4) were removed from our dataset to yield 29 samples.

### 2.9. Statistical analyses

After removing samples with low protein detection, we retained 29 samples for statistical analyses. To identify hotspots vs. low-activity sediments, we examined the distribution of aerobic metabolism rates in all 29 samples (Fig. S4). The distribution was approximately normal with a minimum of 361.83, a mean of 810.49, and a maximum of 1472.86. We defined low-activity sediments as those falling in the bottom quartile of the distribution (maximum rate of aerobic metabolism: 619.45) and high activity samples as those falling in the top quartile of the distribution (minimum rate of aerobic metabolism: 1033.66), yielding a sample size of 7 for each set of samples. All statistical analyses were conducted using R software (https://www.r-project.org/) and graphics were generated with either base packages or ‘ggplot2.’

We used two approaches to determine differences in the composition of metabolites, microbial phylogenies, metagenomic functional potential (i.e., all annotated genes), and metaproteomes between hotspots and low-activity sediments. Metabolomes were analyzed as present/absence data, and all other data types were analyzed as relative abundances. First, we examined differences in full compositional profiles across groups using permutational multivariate analysis of variance (PERMANOVA, 999 permutations, ‘vegan’ package) and accounting for potential non-independence between samples within the same core by stratifiying this analysis by depth (as per Graham et al., 2017b). Because metabolite data were in presence/absence form, a Sorenson dissimilarity distance was constructed and used to compare metabolite data. All other datasets were analyzed using Bray-Curtis dissimilarity to account for differences in relative abundance.

Secondly, we used linear mixed effect models to investigate variation in specific microbial attributes between hotspots and low-activity sediments. Following the convention of Graham et al. (2017b), we included depth as a random effect in these models to account for possible non-independence among sediments. A given microbial attribute was the dependent variable and sediment type was the independent variable. Mixed models were compared to null expectations (i.e., a model including only random effects) with analysis of variance (ANOVA) to determine significance. For significant models (P < 0.05), R^2^ values for microbial attributes independently and for the entire mixed model (i.e., the effects of the attribute plus depth) were determined with the ‘r.squaredGLMM’ function in the ‘MuMIn’ package (Barton, 2009). Because few compounds differed across activity levels, P-values were not adjusted for multiple comparisons in order to maximize the likelihood of detecting differences.

We selected specific attributes for analysis as follows: For metabolomic data, we grouped peaks by their Van Krevlen assignment and compared the relative abundance of each Van Krevlen class across activity groups (e.g., the relative abundance of amino sugars in hotspots vs. low-activity sediments). To identify specific microbial phylogenic and functional groups of interest, we employed the following procedure: (1) we ranked the relative abundance of each assigned phylogeny (class level) or functional annotation (pfam, COG, and KEGG) within each sample; (2) we compiled a master list containing the 25 most abundant phylogenies or functional annotations in every sample; and (3) we determined the relative abundance of every unique phylogeny or functional annotation in the master list within each sample. In this way, we examined every taxon or metabolic pathway that was present in high abundance in at least one sample. For metaproteomic data, we employed a similar pipeline but because the proteomic dataset was smaller, we comprised our master list from any protein in the top 5% of relative abundance in a given sample (equivalent to more than 10 proteins per sample).

Because (1) metagenomes did not vary strongly between hotspots and low-activity sediments, (2) proteins are tightly coupled to function conceptually (i.e., they are proximate catalysts of reactions), and (3) metabolites were dramatically different across activity levels; we focused subsequent analyses only on metaproteomic and metabolomics data. We first investigated the extent to which metaproteome structure corresponded to metabolome structure in the full dataset (n = 29) by comparing dissimilarity in metabolite composition (Sorenson dissimilarity) to differences in proteomic composition (Bray-Curtis dissimilarity, Mantel test using Spearman correlation and stratifying by depth). We also examined the extent to which each data type corresponded to changes in aerobic metabolism by fitting linear and quadratic mixed effects models across the full dataset. As above, depth was included as a random effect in each model. To reduce multidimensional metaproteomic and metabolomic data into vectors for model construction, we extracted the first two principle components of variation within data type (PCA analysis, ‘prcomp’). PC1 and PC2 were predictors in the mixed effects models, and aerobic metabolism was the response variable. Mixed models were compared to null expectations (i.e., models including only random effects) with ANOVA to determine significance. R^2^ values were determined with the ‘r.squaredGLMM’ function in the ‘MuMIn’ package. Since both PC1 and PC2 from metabolomes were significantly related to aerobic metabolism, we also constructed a model that included both PC1 and PC2 as predictors. Lastly, to examine additional explanatory power that could be gained by incorporating bulk C as predictors along with metabolite chemistry, we constructed a final model with PC1, PC2, and percent C. This model was compared to the model with PC1 and PC2 only using ANOVA to determine if percent C significantly improved explanatory power of aerobic metabolism.

Finally, because metabolomes were most distinguishing between hotspots and low-activity sediments and corresponded to rates of aerobic metabolism, we concluded with set of analyses with metabolomic data only. We deciphered specific metabolites that distinguished hotspots from low-activity sediments using ‘random forest’ machine learning algorithm (Liaw and Wiener, 2002, ‘randomForest’ package). Forests were constructed using metabolite peaks that were assigned chemical formulae, removing singletons for computational limitations. For each forest, 1000 trees were constructed with replacement, and error rates converged near zero (Fig. S5). Peak importance and partial dependence in distinguishing high vs. low activity samples were calculated with the ‘imp’ and ‘PartialPlot’ functions, respectively. Peaks that were assigned a ‘Mean Decrease in Accuracy’ of greater than zero were considered to be distinguishing metabolites. This approach yielded 55 peaks distinguishing low activity samples and 272 peaks distinguishing high activity samples.

We also used metabolite transformations to infer biochemical processes associated with aerobic metabolism in hotspots. We identified transformations of interest using the following procedure: (1) we summed the relative abundance of each transformation across all samples (n = 29) and (2) we chose the transformations whose abundance was in the top 20% of all transformations (18 out of 92, mean relative abundance: 1.6% to 14.5%). By doing so, we analyzed transformations that occurred most frequently in our dataset, irrespective of their occurrence within individual samples.

We then compared the relative abundance of specific transformations between hotspots and low-activity sediments using linear mixed effect models using the same procedure as for abundant microbial phyologenies, metabolic pathways, expressed proteins, and metabolites (described above). Because many abundant transformations in hotspots seemed to contain nitrogen (N) while transformations in low-activity sediments did not, we also examined the role in nitrogenous compounds in biochemical transformations in hotspots. We separated transformations into those containing N and N-free transformations (Table S3 and S4) and determined the relative abundance of transformations in each sample that were either nitrogenous or N-free. We then compared the relative abundance of nitrogenous and N-free compounds between hotspots and low-activity sediments using mixed effects models. Because the distribution of transformations within nitrogenous and N-free categories was highly non-Gaussian, we applied a Box-Cox transformation to each data type (’boxCox’ function in ‘car’ package, Sakia, 1992), and subsequently constructed linear mixed effects models. We further separated amino-acid nitrogenous transformations from non-amino acid nitrogenous transformations (hereafter ‘complex N’, Table S5 and S6), applied a Box-Cox transformation, and constructed linear mixed effects models with these data.

## 3. Results

### 3.1. Multi ‘Omic Differences across Rates of Aerobic Metabolism

Microbiome structure, functional potential, and protein expression showed limited differences between hotspots vs. low-activity sediments. No significant difference was observed in the composition of microbiome phylogenies (P = 0.57, Fig. 1a), metagenomic annotations (P = 0.69, Fig. 1b), and metaproteomes (P = 0.91, Fig. 1c) between hotspots and low-activity sediments (PERMANOVA). As well, major clades, pathways, and proteins were similar among sediments (Fig. 1).

**Fig. 1.**
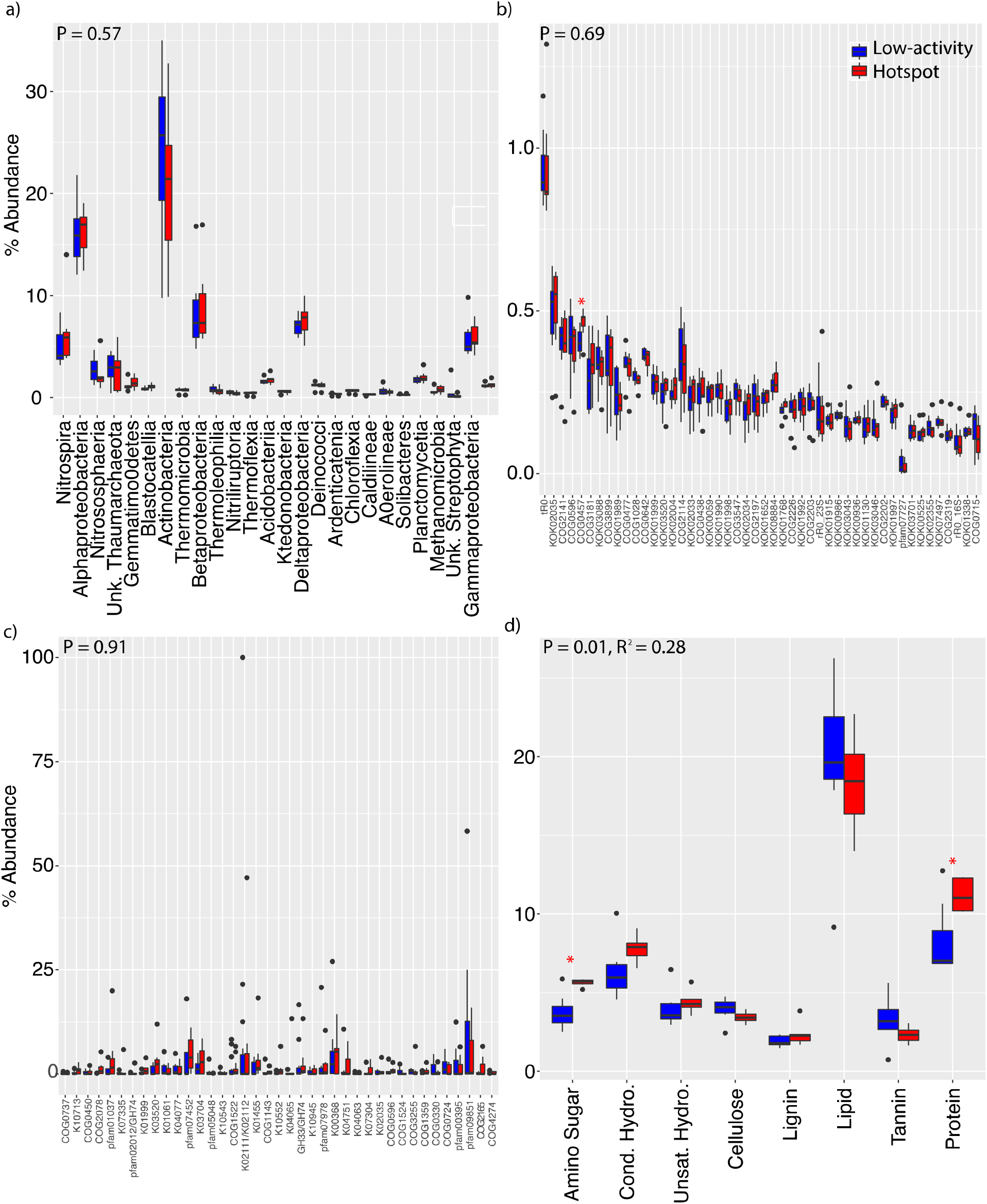
Multi ‘Omic Differences in Hotspots vs. Low-activity Sediments. (a-d) show the most abundant classifications only for visual simplicity, but P-values in the upper left-hand corner of each panel are derived from all data. P-values are derived from PERMANOVA with stratification by depth. R^2^ values are provided for significant P-values. (a) shows abundant microorganisms (grouped at the class level), (b) shows abundant metagenomics annotations, (c) shows abundant metaproteomic identifications, and (d) shows metabolomics data grouped by compound class. Asterisks denote significant differences between hotspots and low-activity sediments, via mixed models with depth as a random effect (P < 0.05).

Metabolome composition, however, differed between hotspots and low-activity samples (P = 0.01, R^2^ = 0.28, Fig. 1d, PERMANOVA). When grouped into major compound classes, hotspots contained more amino sugar- and protein-like metabolites (P < 0.05, Fig. 1d, mixed effects). Metabolite pool composition was uncorrelated to proteome structure (P = 0.62, Fig. S6, Mantel test across all samples).

Finally, when considering aerobic metabolism as a continuous variable, metabolome composition was tightly correlated to aerobic metabolism. The first two principle coordinates of metabolite pool composition had significant independent relationships with aerobic metabolism and together explained 67% of variation in aerobic metabolism (mixed effects, P (PC1, quad.) = 0.0006 & P (PC2, linear) = 0.01, P (combined, linear) << 0.0001, Fig. 2a, Fig. S7). Statistical models including both bulk C and metabolite composition were not significantly different from models including metabolite composition alone (R^2^ 0.73 vs. 0.67, P = 1). Metaproteome composition was uncorrelated to aerobic metabolism despite representing the proximal catalysts of metabolic processes (mixed effects, P (PC1) = 0.29 & P (PC2) = 0.53, Fig. 2b).

**Fig. 2.**
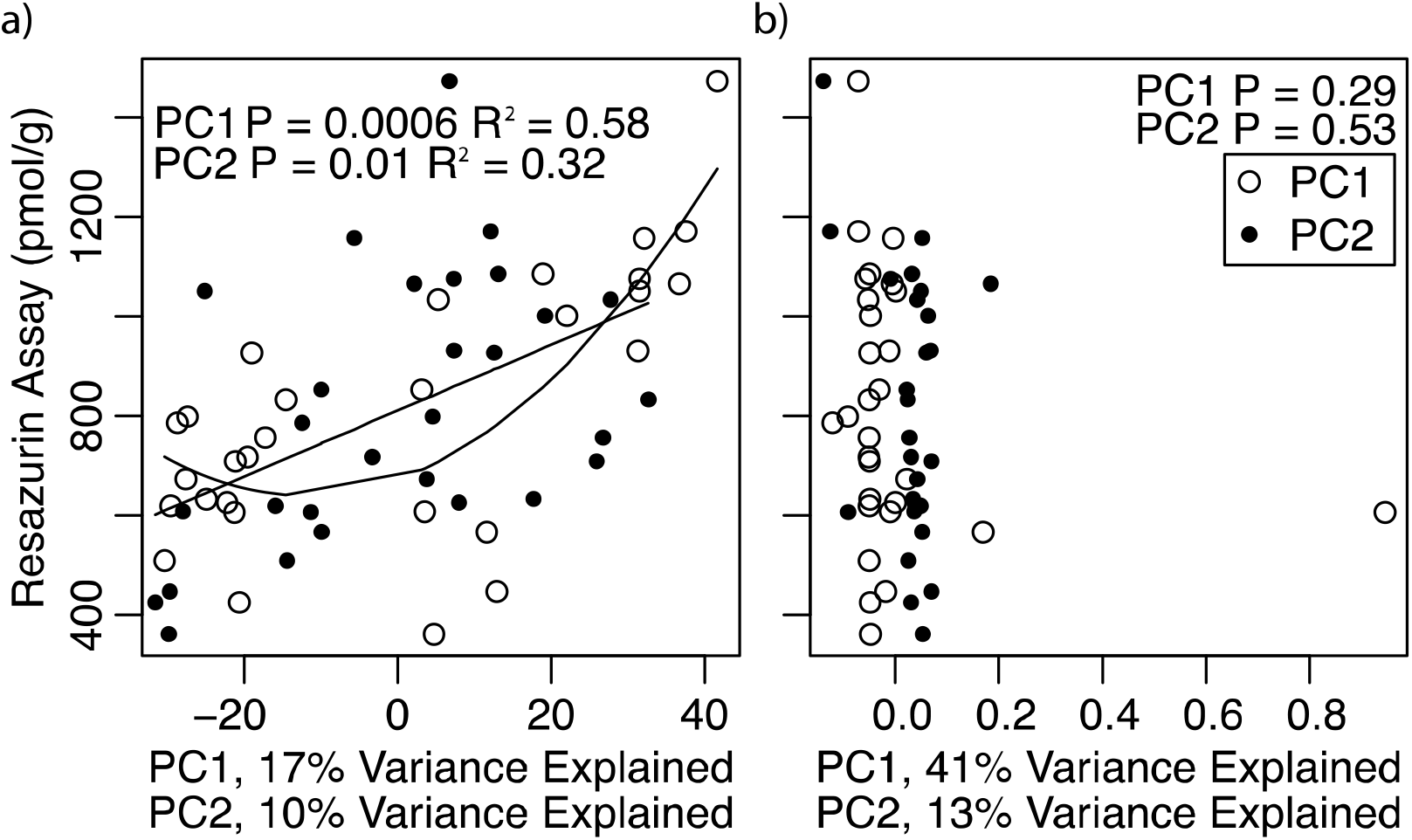
Relationships of Metabolomes and Metaproteomes with Aerobic Metabolism. The first two principle components were extracted from metabolomics and metaproteome data. Correlations between each component and aerobic metabolism were determined using linear and quadratic mixed models that included depth as a random effects. Regression lines are shown for the best model; only significant models are shown. (a) Metabolomic data were strongly correlated with aerobic metabolism, while (b) metaproteomic data were uncorrelated.

### 3.2. Distinguishing Metabolite Chemistry and Transformations

Because hotspots were discriminated from low-activity sediments only in the metabolome, we examined metabolite pools to identify compounds and infer processes that distinguished hotspots from low-activity sediments. Hotspots were characterized by high molecular weight (P < 0.0001, Fig. 3a), chemically diverse (Fig. 3b) compounds. In particular, nitrogenous classes of compounds (i.e., amino sugars and proteins) were present in metabolites that described hotpots and absent from metabolites characterizing low activity sediments (Fig. 3b).

**Fig. 3.**
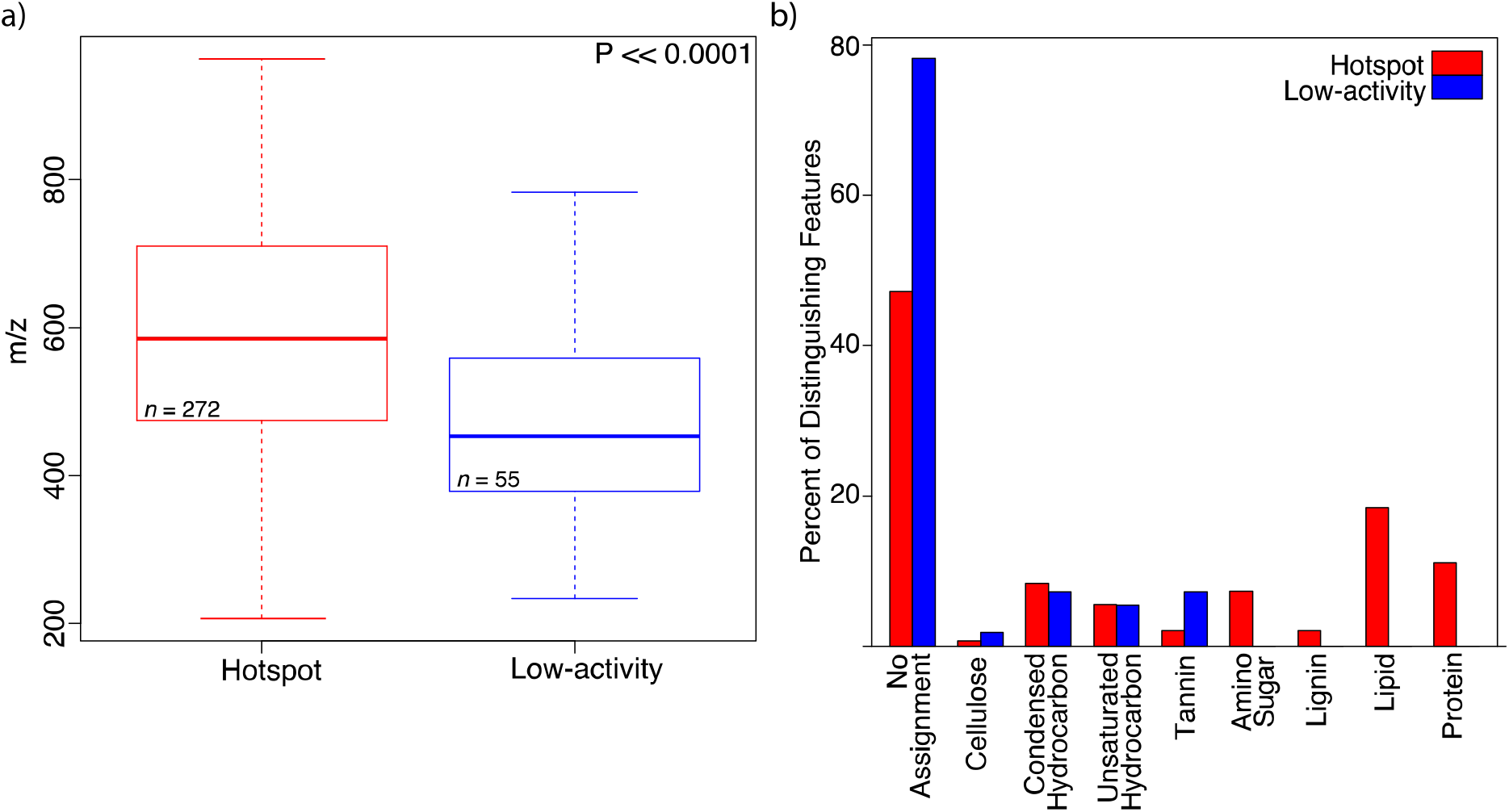
Metabolites Distinguishing Hotspots from Low-Activity Sediments. (a) The molecular weight of distinguishing metabolites was significantly higher in hotspots than low-activity sediments. A boxplot of molecular weights are shown, with the center line indicating the mean and the hinges of each box representing the values at the 25^th^ and 75^th^ percentiles. The whiskers represent the range of the data, calculated using default settings in R. P-value is derived from a one-sided Mann-Whitney U test. (b) shows the distribution of distinguishing metabolites in hotspots vs. low-activity sediments. Compounds are grouped by molecular class and displayed as a percent of distinguishing metabolites in each class.

Hotspots also exhibited a more even distribution of abundant biochemical transformations, indicating greater diversity of biogeochemical processes in these sediments relative to low-activity sediments (Fig. 4a). These transformations were inferred by comparing differences between peaks in FT-ICR-MS data to a database of known biochemical transformations (see section 2.6). Further, transformations involving nitrogenous compounds (P = 0.03 R^2^ = 0.26, Table S3), particularly those involving complex N (P = 0.04 R^2^ = 0.20, Fig. 4b, Table S6)), were more prevalent in hotspots.

**Fig. 4.**
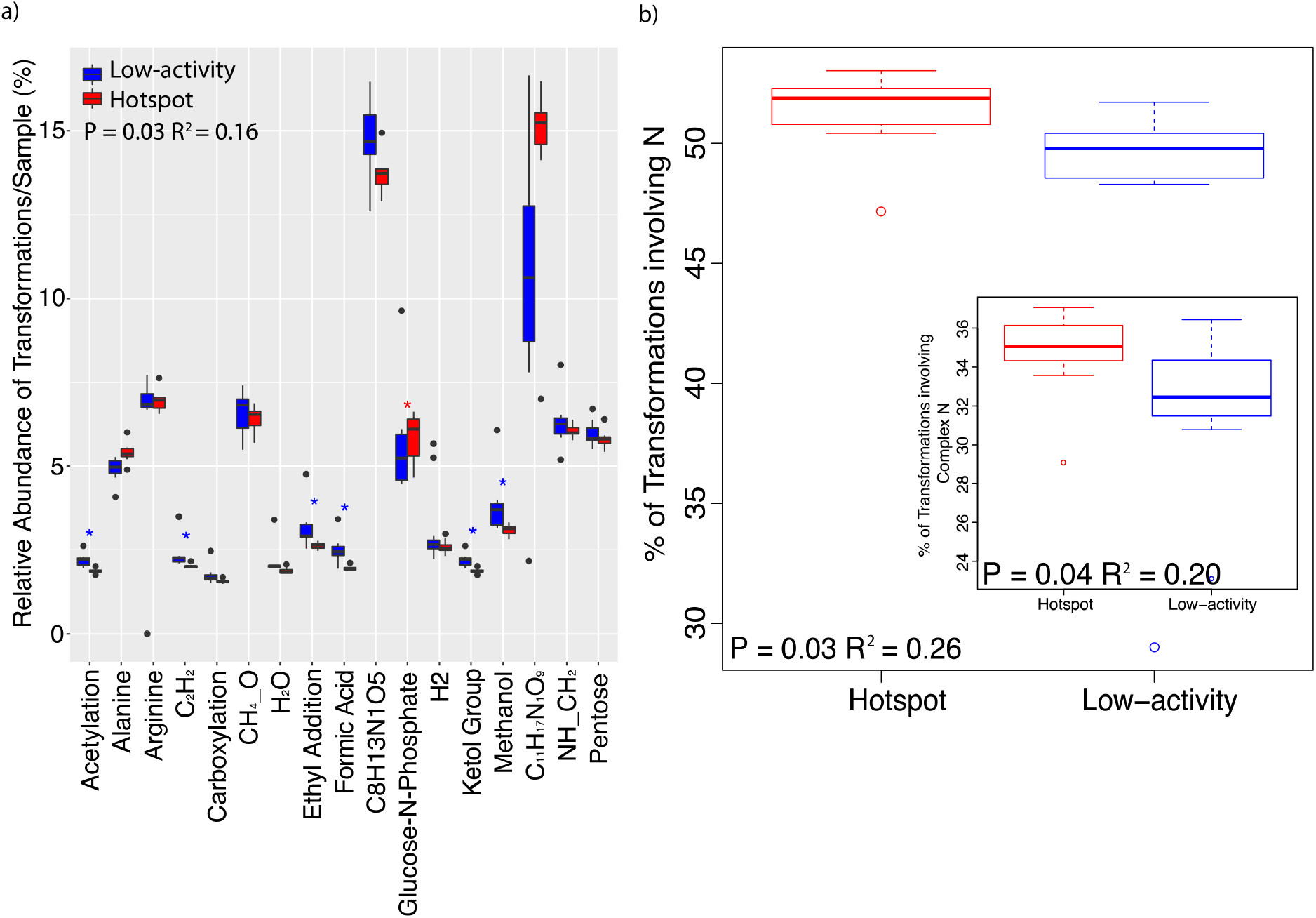
Differences in Biochemical Transformations between Hotspots and Low-activity Sediments. (a) shows the most abundant transformations only for visual simplicity, but the P-value in the upper left-hand corner is derived from all data. Asterisks denote significant differences between hotspots and low-activity sediments, via mixed models with depth as a random effect (P < 0.05). (b) shows a boxplot of the percent of transformations involving N across hotspots and low-activity sediments. The inset shows the same analysis using only transformations involving ‘complex N’. Lists of the specific transformations included in each analysis are shown in Tables S3-S6. For each boxplot, with the center line indicating the mean and the hinges of each box representing the values at the 25^th^ and 75^th^ percentiles. The whiskers represent the range of the data, calculated using default settings in R. P-values and R^2^ are calculated using mixed models with depth as a random effect. Due to non-normality in the distribution of transformations, a Box-Cox transformation was applied prior to model construction.

## 4. Discussion

Our multi ‘omics investigation of microbiome structure and function in hyporheic zone hotspots highlights the importance of vegetation in driving local-scale (<200m^2^) aerobic metabolism in the subsurface. Pronounced spatial heterogeneity in metabolism across our relatively small spatial domain (∼175 x 10m) is inconsistent with representations of biogeochemical processes within ecosystem models that often aggregate environmental properties across similarly sized domains, and suggests a need for finer scale representation of processes in these models. Our research supports previous observations of relationships between riparian vegetation and hyporheic zone hotspots due to associated carbon and nutrient inputs (Harms and Grimm, 2008; McClain et al., 2003; Schade et al., 2001). For example, Kuglerova et al. (2014) demonstrated that increased vegetation growth corresponded to enhanced biogeochemical cycling in areas of the hyporheic zone experiencing maximal groundwater-surface water mixing. In our system, hotspots were disproportionately distributed within sediments underlying vegetation and exhibited up to 4 times the rate of aerobic metabolism in low-activity sediments despite a small sampling domain (Fig. S4).

Contrary to our hypothesis that microbiome composition (metagenomics) and expression (metaproteomics) would be shifted towards carbohydrate metabolism in hotspots, our results support a predominant role for phenotypic plasticity within sediment microbiomes. Phenotypic plasticity, whereby an organism encodes multiple metabolic pathways and adjusts the expression of each depending on environmental conditions, is common in free-living microbes and manifests as different metabolisms occurring in samples with the same genomic composition. Microbiome phylogeny and functional potential did not vary with the level of aerobic respiration in our sediments. While Hartman *et al*. (2017) demonstrated a consistent metagenomic shift from aromatic to carbohydrate metabolism with increased carbon turnover in global soils, we observed no such relationship in our sediments. A key difference between Hartman *et al*. (2017) and the present study is that Hartman *et al*. demonstrated decreased nutrient uptake with increased activity. In the present study, decreased nutrient uptake would correspond to fewer nitrogenous biochemical transformations, whereas we observed increasing activity of biochemical pathways involving nitrogen in hotspots vs. low-activity sediments. The lack of microbiome change in our study may therefore be constrained by N-limitation. For instance, if microbiomes in our sediments are always under strong selection for N-harvesting metabolisms, they would be structured to contain organisms with the ability to metabolize organic N (Craine et al., 2007; Moorhead and Sinsabaugh, 2006), regardless of the magnitude of available N.

Interestingly, while one would expect protein expression to correlate with changes in activity, we saw no such effect in our metaproteomic data. The long residence time of enzymes compared to hydrologic mixing may decouple expression from function in dynamic environments such as the hyporheic zone. Hydrologic exchange in the hyporheic zone occurs on relatively short time scales (commonly minutes to days), as well as longer seasonal exchanges, and creates fleeting periods of nutrient availability (Cardenas et al., 2008; Fritz and Arntzen, 2007; Sawyer and Cardenas, 2009; Zarnetske et al., 2011). Recent work in soils has shown that protein decay rates vary widely across enzymes and that some enzymes can persist more than 12 weeks in soils (Schimel et al., 2017). Assuming a similar magnitude of decay in hyporheic zone sediments would generate a disconnect between protein expression and resource availability that may ultimately disassociate enzyme pool composition from respiration rates, leading to the lack of relationship between metaproteome composition and aerobic metabolism observed here.

Our results indicate that improved resolution into sediment metabolomes provides greater insight into sediment function than bulk C content at a given point in time. While there is a long history of constraining biogeochemical rates with standing resource pools, our work indicates that bulk C alone was an insufficient predictor of aerobic respiration (Fig. S8). Metabolite chemistry explained 67% of variation in aerobic metabolism (in comparison to 42% by bulk C pools, Fig. S8), and including bulk C content as a predictor in addition to metabolome composition did not improve the explanatory power of our statistical models. In particular, nitrogenous metabolites were associated with enhanced metabolism. We therefore suggest that metabolome composition can explain aerobic metabolism better than bulk C stocks in some systems and subsequently that new process-based model structures need to account for multiple C pools with various organic N content. For instance, current generation models incorporate mechanisms reflecting that certain C molecules (e.g., glucose) can enter microbial cells by passing directly through their cellular membranes, while larger molecules require secreted extracellular enzymes to break them down prior to microbial uptake (Wang et al., 2013; Wieder et al., 2014). Extending these new frameworks to further parse C chemistry by organic N content and associated metabolic pathways could advance the performance of these models.

Microbiomes associated with hyporheic zone hotspots metabolized a broader range of resources compared to microbiomes in low-activity sediments. Enhanced biogeochemical cycling can be achieved through microorganisms using different portions of resource pools (termed ‘resource partitioning’ or ‘niche complementarity’) in which diverse microorganisms use different portions of resource pools (Cardinale, 2011; Loreau and Hector, 2001; Schoener, 1974). Consistent with this mechanism, we show a divergence in active metabolic pathways (inferred from biochemical transformations) in hotspots compared to low-activity sediments. Thus, despite similar microbiome genomic content, hotspots seem to result from the expansion of realized niches of extant microbiomes. In this case, increased biogeochemical function arises through phenotypic plasticity.

Hotspots contained diverse metabolites, were distinguished from low-activity sediments by protein- and amino sugar-like metabolites, and exhibited higher instances of biochemical transformations involving organic N. It has been widely demonstrated that N availability can limit microbial activity (Treseder, 2008), resulting in slower decomposition and ecosystem organic matter processing (Averill and Waring, 2017; Weintraub and Schimel, 2003; Zhang et al., 2008); and previous work in this system has suggested a coupling of C and N cycles (Stegen et al., 2018). An excess of inorganic N relative to other N sources has also been demonstrated to shift microbial metabolism by suppressing oxidative enzyme synthesis (Edwards et al., 2011; Jian et al., 2016) and by changing the ratios of synthesized enzymes that degrade recalcitrant vs. labile organic matter (Moorhead and Sinsabaugh, 2006). Similarly our analysis of metabolite transformations supports different metabolic strategies among hotspot vs. low-activity microbiomes as well as a role for organic N in particular as a driver of aerobic metabolism in hotspots.

While C:N ratios do not reflect obvious N limitation in our system (means: 6.55 (hotspot) and 6.19 (low-activity)), these ratios can misrepresent chemical conditions experienced by microbiomes due to N absorption to soil matrices and by aggregation processes that physically separate N from microorganisms (Lützow et al., 2006; Sollins et al., 1996). For instance, Darrouzet-Nardi and Weintraub (2014) demonstrated that bulk methods of C and N determination such as those used here may overestimate N availability by 5-fold. As such, organic N may play a critical role in our system despite ratios of C:N in bulk sediments.

Moorhead and Sinsabaugh (2006) proposed a concept known as ‘microbial nitrogen mining’ in which microorganisms in environments with limited availability of labile N metabolize C to access organic N (Craine et al., 2007; Moorhead and Sinsabaugh, 2006). Microbial N mining should result in the preferential decomposition of organic N, and our results indicate that this process may contribute to high respiration rates in hyporheic zone hotspots. Consistent with N mining, we observed biochemical pathways involving nitrogenous compounds, and complex N in particular within hotspots, supporting previous observations that organic N increases the decomposition of organic matter (reviewed in Averill and Waring, 2017; Hu et al., 2001; Zhang et al., 2008). The metabolism of complex nitrogenous organic compounds specifically suggests that the breakdown of more chemically-complex organic material may be stimulated to facilitate microbial access to nitrogen and that more available labile C in vegetated sediments provides additional energy necessary to metabolize more chemically-complex nitrogenous molecules.

Based on our results, we propose a new conceptualization of hyporheic zone organic matter transformations in which greater resolution into C metabolites and associated metabolic pathways––beyond bulk C stocks and/or microbiome structure––is critical for predictive hydrobiogeochemical models (Fig. 5). Phenotypic plasticity suggests (1) an ability for microbiomes to rapidly acclimate to environmental conditions, regardless of their underlying genomic content, and (2) that there may be limited contributions of metagenomic information to functional predictions in dynamic systems. Because compositional shifts in microbiomes are often on the order of days to months or longer (Balser and Firestone, 2005; Graham et al., 2016a; Poretsky et al., 2014; Waldrop and Firestone, 2006), the ability of microorganisms to adjust metabolic function to suit prevailing conditions without associated species turnover denotes that the timescale of microbiome response is not limited to these longer timescales and instead that microbial responses in some systems may be more tightly constrained by RNA transcription and translation (∼minutes or less, Moran et al., 2013).

**Fig. 5.**
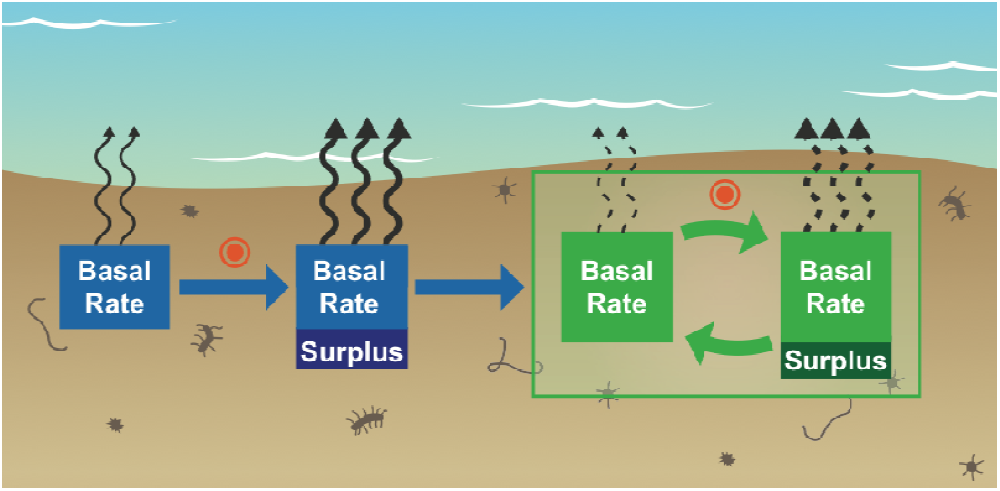
Conceptualization of a Role for Metabolic Pathways in CO_2_ Flux Predictions. We propose that greater resolution into C metabolites and associated metabolic pathways––beyond sediment bulk C and/or microbiome structure––is vital for predictive hydrobiogeochemical models. Broader metabolic activity in biogeochemical hotspots stimulates the production and consumption of diverse metabolites, in particular nitrogenous compounds. Therefore, there is a need to incorporate subsequent feedbacks between metabolite consumption and production into predictive biogeochemical models. Orange bull’s eyes represent the occurrence of a hotspot or hot moment. Basal rate is the baseline aerobic metabolism of sediments. Surplus metabolism indicates the increase in aerobic metabolism associated with hotspots and includes stimulated organic N respiration. Arrows denote CO_2_ flux, where the number and thickness of arrow denotes the relative size of the flux. Blue boxes indicate the initial condition of sediments and the first occurrence of a hotspot or hot moment. Green boxes show an ongoing feedback loop in which the basal rate of a microbiome is continuously influenced by metabolite production and consumption.

Further, we suggest that broader metabolic activity in biogeochemical hotspots stimulates the production and consumption of diverse metabolites which alter the composition of future resource pools, and therefore that understanding subsequent feedbacks between metabolite consumption and production is vital for predictive biogeochemical models. For example, as metabolism proceeds through time, future metabolic rates are dependent on the resources (i.e., metabolites) generated by these pathways. Metabolism of an expanded suite of resources through broader realized niche space within biogeochemical hotspots may therefore fundamentally change the subsequent metabolite and resource profiles (Fig. 5). As hotspots and hot moments occur through time and space, resource pools progressively change in ways that are dependent on microbial transformations of prior resource pools, and knowledge about the specific metabolic pathways that generate propagating changes as well as their rates and limitations is imperative for improving biogeochemical predictions. Our results also demonstrate organic N in particular may be involved these alternative pathways, and thus, that nutrient limitations are important considerations in constraining rates within predictive models, as has been recently explored by Wieder et al (2015b). We further extended this conceptualization to propose that organic N specifically should be considered as a possible regulatory factor in process-based ecosystem models.

The fate of organic carbon stored in soils and sediments, particularly at terrestrial-aquatic interfaces (Battin et al., 2009; Boulton et al., 1998; Cole et al., 2007; Marín-Spiotta et al., 2014; Regnier et al., 2013), is responsible for large amounts of uncertainty in predictions of future global biogeochemistry (Luo et al., 2016; Todd-Brown et al., 2013). Here, we provide insight into the molecular mechanisms generating hotspots of elevated aerobic metabolism in the hyporheic zone. While microbiome structure and protein expression did not vary across levels of aerobic metabolism, we found that metabolite chemistry and diverse, N-related biochemical pathways were associated with hyporheic zone hotspots. We posit that microbiome phenotypic plasticity can enable microbiomes with similar structure to expand their realized niche in response to changing environments and rapidly increase function in favorable environments. Because of fleeting environmental changes within hyporheic zones, we hypothesize that hyporheic zone microbiomes may exhibit more phenotypic plasticity than more stable ecosystems, meriting further investigation as we explore spatiotemporal dynamics across multiple systems (Fodelianakis et al., 2017). As we search for consensus on the importance of various environmental factors in predicting future biogeochemical rates (Graham et al., 2016b; Graham et al., 2014; Luo et al., 2016; Wieder et al., 2017), a spatially-explicit understanding of the mechanisms regulating organic matter transformations in areas with elevated biogeochemical rates is essential for improving the conceptualization and parameterization of ecosystem models. We propose that understanding and constraining the rates of specific metabolic pathways that utilize discrete portions of resource pools is a critical consideration when advancing carbon cycle complexity in these models.

## Acknowledgements

This research was supported by the US Department of Energy (DOE), Office of Biological and Environmental Research (BER), as part of Subsurface Biogeochemical Research Program’s Scientific Focus Area (SFA) at the Pacific Northwest National Laboratory (PNNL). PNNL is operated for DOE by Battelle under contract DE-AC06-76RLO 1830. Data were generated under JGI/EMSL user proposal 1781. A portion of the research was performed at Environmental Molecular Science Laboratory User Facility. All data are publicly available at DOI XXXXXXX.

