## Supplementary Materials for "Multi ‘omics comparison reveals metabolome biochemistry, not microbiome composition or gene expression, corresponds to elevated biogeochemical function in the hyporheic zone"

**SUPPLEMENATL MATERIAL**

*1.1 Extended Description of Physicochemical Assays*

1.1.1 Particle Size Distribution.

Samples were dried for 4-5 days at 105°OC, weighed, then transferred to 4L polypropylene Nalgene bottles and vigorously agitated to break up aggregates. Material was then sieved into1- phi size classes from 64mm (-6 phi) to <.062mm (5 phi) and weighed to determine the weight percentage for each fraction. Each subsample was placed in a 50ml Corning tube with 25ml of 1% pyrophosphate solution and placed on a rotary shaker overnight. This mixture was then poured and washed through a #230 sieve (.062mm). Material retained on the sieve was dried overnight and weighed to determine the weight percentage of the sand fraction, which was then subtracted from the initial 10g to determine the silt and clay fraction. These measurements were then averaged and the weight-by-size fractions of the original 10g were determined. We used the Modified Wentworth Scale to define the mud fraction as <.062mm (#230 sieve), the sand fraction as <2mm to .062 (passes #10 sieve, but retained on a #230), and the gravel fraction as >2mm (retained on #10 sieve) to 64mm.

1.1.2 Sample Processing.

Samples were transferred to an anaerobic glovebag with 95% N_2_ and 5% H_2_ (Coy Laboratory Products, Grass Lake, MI) and thawed on clean 2mm stainless steel sieves prior to processing. Samples were sieved and homogenized; and 20g was subsampled into a pre-weighed, pre-cleaned 40ml glass vial for Fe(II) analysis. The remaining <2mm sieved and >2mm bulk samples were removed from the glovebag. Subsamples were collected from the <2mm fraction for further analysis. Approximately 5g was transferred into a 40-mL borosilicate glass vial and stored at -80^o^C for analysis of OC chemistry (see below). An additional sample was taken for elemental analysis (OC, N) and remaining material was divided into 20g samples collected into 40-mL borosilicate glass vials and stored at -80^o^C.

1.1.3. Analytical methods.

We analyzed sample geochemistry with the following approaches. Total carbon (OC), nitrogen (N), and sulfur (S) were measured using an Elementar vario EL cube (Elementar Co.Germany). To analyze ammonia (NH_4_^+^), we extracted 2g sediment with 4mL 2M KCl for 1hr on a reciprocal shaker. Extracts were filtered (0.22 um) and NH_4_ was measured with Hach Kit 2604545 (Hach, Loveland, Co). Fe(II) content was assessed by ferrozine assay[*Stookey*, 1970]. Briefly, 1g sediment was extracted for 1hr with 10 mL 0.5N HCl with 200 rpm shaking in an anaerobic chamber and filtered through 0.22um PTFE syringe filters. One mL of filtrate and 1 mL Ferrozine were combined in a 1.5 mL, and fluorescence was measured at 562 nm to determine Fe(II) content. Total Fe was determined by inductively coupled plasma (ICP), and Fe(III) was calculated as the difference between Fe(II) and total Fe. All other ion concentrations were measured by ICP on 0.5N HCl extractions, as described above.

In addition, we assessed aerobic metabolism in each sample with a resazurin reduction assay, modified from Haggerty *et al.*[*Haggerty et al.*, 2009]. We added 6ml of filtered river water from the Columbia River to four replicate 1 cc subsamples, taken with a cut-off syringe, from each depth. One replicate vial from each location was heat killed in a 90^o^C water bath for 1 hour, and cooled on ice to bring back to 4^o^C. Resazurin incubations were started by adding 200μl of 30μM resazurin to vials when cold, gently mixing and then incubating on an angle at 50rpm and 21^o^C. After 4hr, vials were weighed and 6mL acetonitrile (ACN) added to begin a 1hr extraction. After ACN addition, vials were sealed, vortexed and weighed again before being placed in a sonicator bath for 10min. After sonication, vials were put back on the 50rpm shaker. After a 1hr extraction, vials were vortexed and sand was allowed to settle. The screw cap was removed to allow extract to be drawn into a 5-mL syringe fitted with a 20G needle. Extracts were filtered with 33mm, 0.2μm syringe filters (PES, Millex by Millipore) into pre-labeled 12-mL amber vials (Thermoscientific) and stored at 4^o^C. The vial with the remaining sand was dried in a convection oven at 75 ^o^C for at least 72 hrs then weighed. Fluorescence emission maxima for resazurin (630nm) and resorufin (585nm) were measured on resazurin sample extracts using a Horiba Fluorolog 3 fluorimeter. 2mL extract was added to 0.2mL 100mM HEPES (pH 8) in quartz cuvettes and fluorescence intensity quantified by comparison to resazurin and resorufin standards made up in ACN:H_2_O (1:1). Acetonitrile extraction was found to greatly enhance recovery of resazurin and resorufin in preliminary experiments where about 80% of added resazurin was adsorbed to abiotic sand treatments. Typical recoveries with acetonitrile extraction were above 90%. Results of resazurin reduction were expressed in pmoles of resazurin reduced to resorufin per gram dry weight of sand over the 48hr incubation period.

*1.2 Extended Description of FT-ICR-MS Procedures*

Sediment extracts were prepared by adding 1 ml of solvent to 100 mg bulk sediment and shaking in 2 mL capped glass vials for two hours on an Eppendorf Thermomixer. Samples were removed from the shaker and left to stand before spinning down and pulling off the supernatant to stop the extraction. The residual sediment was dried with nitrogen gas to remove any remaining solvent, and then the next solvent was added. The CHCl_3_ and H_2_O extracts were diluted in MeOH to improve ESI efficiency. Tfaily et al. (2015) estimated the extraction efficiency to be ~15%. Tfaily et al. (2015) have previously demonstrated extraction efficiencies as low as 2% to be representative of metabolite pool composition. We further note that numerous studies have established FT-ICR-MS as a robust method for distinguishing compositional differences among carbon metabolites (Herzsprung et al., 2017; Kellerman et al., 2015; Rossel et al., 2016; Ward and Cory, 2015; Zhang et al., 2016).

Ultra-high resolution mass spectrometry of the three different extracts from each sample was carried out using a 12 Tesla Bruker SolariX FT-ICR-MS located at the Environmental Molecular Sciences Laboratory (EMSL) in Richland, WA, USA. As per Tfaily et *al.* (2017), we performed weekly calibration using a tuning solution containing C_2_F_3_O2, C_6_HF_9_N_3_O, C_12_HF_21_N_3_O, C_20_H_18_F_27_N_3_O_8_P_3_, and C_26_H_18_F_39_N_3_O_8_P_3_ with m/z ranging from 112 to 1333 (Agilent Technologies, Santa Clara, CA USA), and instrument settings were optimized using Suwannee River Fulvic Acid (IHSS). The instrument was flushed between samples using a mixture of water and methanol. Blanks were analyzed at the beginning and the end of the day to monitor for background contaminants.

The extracts were injected directly into the mass spectrometer and the ion accumulation time was optimized for all samples to account for differences in carbon concentration. The ion accumulation time ranged between 0.5 and 1s. A standard Bruker electrospray ionization (ESI) source was used to generate negatively charged molecular ions. Samples were introduced to the ESI source equipped with a fused silica tube (30 μm i.d.) through an Agilent 1200 series pump (Agilent Technologies) at a flow rate of 3.0 μL min^-1^. Experimental conditions were as follows: needle voltage, +4.4 kV; Q1 set to 50 *m*/*z*; and the heated resistively coated glass capillary operated at 180 °C.

*1.3 Extended Description of FASP Digestion for Metaproteomics*

Up to 100µl of SDS-Tris buffer (4% SDS, 100mM DTT in 100 mM Tris-HCl, pH 8.0) was added to each pellet and gently sonicated into solution. The protein solutions were transferred into 1.5 mL microcentrifuge tubes and incubated at 95°C for 5 mins and allowed to cool at 4°C for 10 mins. The samples were centrifuged at 15,000x g for 10 minutes to pellet debris and transfered into 30 K molecular weight cut off (MWCO) 500 µl spin filter provided in Expedeon FASP Kits (Expedeon, San Diego, CA), along with 400µl of 8M urea solution. The spin filters were centrifuged at 14,000x g for ~30 mins until the samples hit the dead volume. The waste was removed from the collection tubes and 400 µl of 8M urea solution was added to each sample and centrifuge as described previously and repeated 1 more time for a total of 3 urea spins. 400µl of 25 mM ammonium bicarbonate, pH 8 (ABC) was added and centrifuged as described and repeated for a total of 2 ABC washes. The spin columns were transferred into fresh labeled collection tubes (gently wiping any excess urea off the spin filter). 75 µl of ABC was added to the spin filter along with 4 µl of 1 µg/µl trypsin (affymetrix, Santa Clara, CA) and vortexed. The samples were incubated at 37°C for 3 hrs in a thermomixer with a thermotop to reduce condensation (Eppendorf, Hamburg, Germany). After digestion, 40ul of ABC was added to the sample and centrifuged at 14,000x g for 20 mins. Another 40ul of ABC was added to the top of the spin filter, vortexed and centrifuged again for 10 mins. The spin filter was discarded and the collected peptides were dried in a speed vac to a volume of ~30 µl. A final bicinchoninic acid (BCA) assay (Thermo Scientific, Waltham, MA) was performed to determine the peptide concentration and vialed for MS analysis.

*1.4 Downloadable List of Metaproteomics Files Combined for MS/MS analysis*

<https://genome.jgi.doe.gov/GWRWS3_50_60_FD/GWRWS3_50_60_FD.download.html>

<https://genome.jgi.doe.gov/GWRWS3_40_50_FD/GWRWS3_40_50_FD.download.html>

<https://genome.jgi.doe.gov/GWRWS3_30_40_FD/GWRWS3_30_40_FD.download.html>

<https://genome.jgi.doe.gov/GWRWS3_20_30_FD/GWRWS3_20_30_FD.download.html>

<https://genome.jgi.doe.gov/GWRWS3_10_20_FD/GWRWS3_10_20_FD.download.html>

<https://genome.jgi.doe.gov/GWRWS2_50_60_FD/GWRWS2_50_60_FD.download.html>

<https://genome.jgi.doe.gov/GWRWS2_40_50_FD/GWRWS2_40_50_FD.download.html>

<https://genome.jgi.doe.gov/GWRWS2_30_40_FD/GWRWS2_30_40_FD.download.html>

<https://genome.jgi.doe.gov/GWRWS2_20_30_FD/GWRWS2_20_30_FD.download.html>

<https://genome.jgi.doe.gov/GWRWS2_10_20_FD/GWRWS2_10_20_FD.download.html>

<https://genome.jgi.doe.gov/GWRWS1_50_60_FD/GWRWS1_50_60_FD.download.html>

<https://genome.jgi.doe.gov/GWRWS1_40_50_FD/GWRWS1_40_50_FD.download.html>

<https://genome.jgi.doe.gov/GWRWS1_30_40_FD/GWRWS1_30_40_FD.download.html>

<https://genome.jgi.doe.gov/GWRWS1_30_40_FD/GWRWS1_30_40_FD.download.html>

<https://genome.jgi.doe.gov/GWRWS1_10_20_FD/GWRWS1_10_20_FD.download.html>

<https://genome.jgi.doe.gov/GWRWS3_0_10_FD/GWRWS3_0_10_FD.download.html>

<https://genome.jgi.doe.gov/GWRWS2_0_10_FD/GWRWS2_0_10_FD.download.html>

<https://genome.jgi.doe.gov/GWRWS1_0_10_FD/GWRWS1_0_10_FD.download.html>

<https://genome.jgi.doe.gov/GWRWN3_50_60_FD/GWRWN3_50_60_FD.download.html>

<https://genome.jgi.doe.gov/GWRWN3_40_50_FD/GWRWN3_40_50_FD.download.html>

<https://genome.jgi.doe.gov/GWRWN3_30_40_FD/GWRWN3_30_40_FD.download.html>

<https://genome.jgi.doe.gov/GWRWN3_20_30_FD/GWRWN3_20_30_FD.download.html>

<https://genome.jgi.doe.gov/GWRWN3_10_20_FD/GWRWN3_10_20_FD.download.html>

<https://genome.jgi.doe.gov/GWRWN2_50_60_FD/GWRWN2_50_60_FD.download.html>

<https://genome.jgi.doe.gov/GWRWN2_40_50_FD/GWRWN2_40_50_FD.download.html>

<https://genome.jgi.doe.gov/GWRWN2_30_40_FD/GWRWN2_30_40_FD.download.html>

<https://genome.jgi.doe.gov/GWRWN1_40_50_FD/GWRWN1_40_50_FD.download.html>

<https://genome.jgi.doe.gov/GWRWN1_30_40_FD/GWRWN1_30_40_FD.download.html>

<https://genome.jgi.doe.gov/GWRWN1_20_30_FD/GWRWN1_20_30_FD.download.html>

<https://genome.jgi.doe.gov/GWRWN1_10_20_FD/GWRWN1_10_20_FD.download.html>

<https://genome.jgi.doe.gov/GWRWN3_0_10_FD/GWRWN3_0_10_FD.download.html>

<https://genome.jgi.doe.gov/GWRWN2_0_30_FD/GWRWN2_0_30_FD.download.html>

<https://genome.jgi.doe.gov/GWRWN1_0_10_FD/GWRWN1_0_10_FD.download.html>

**Figures.**

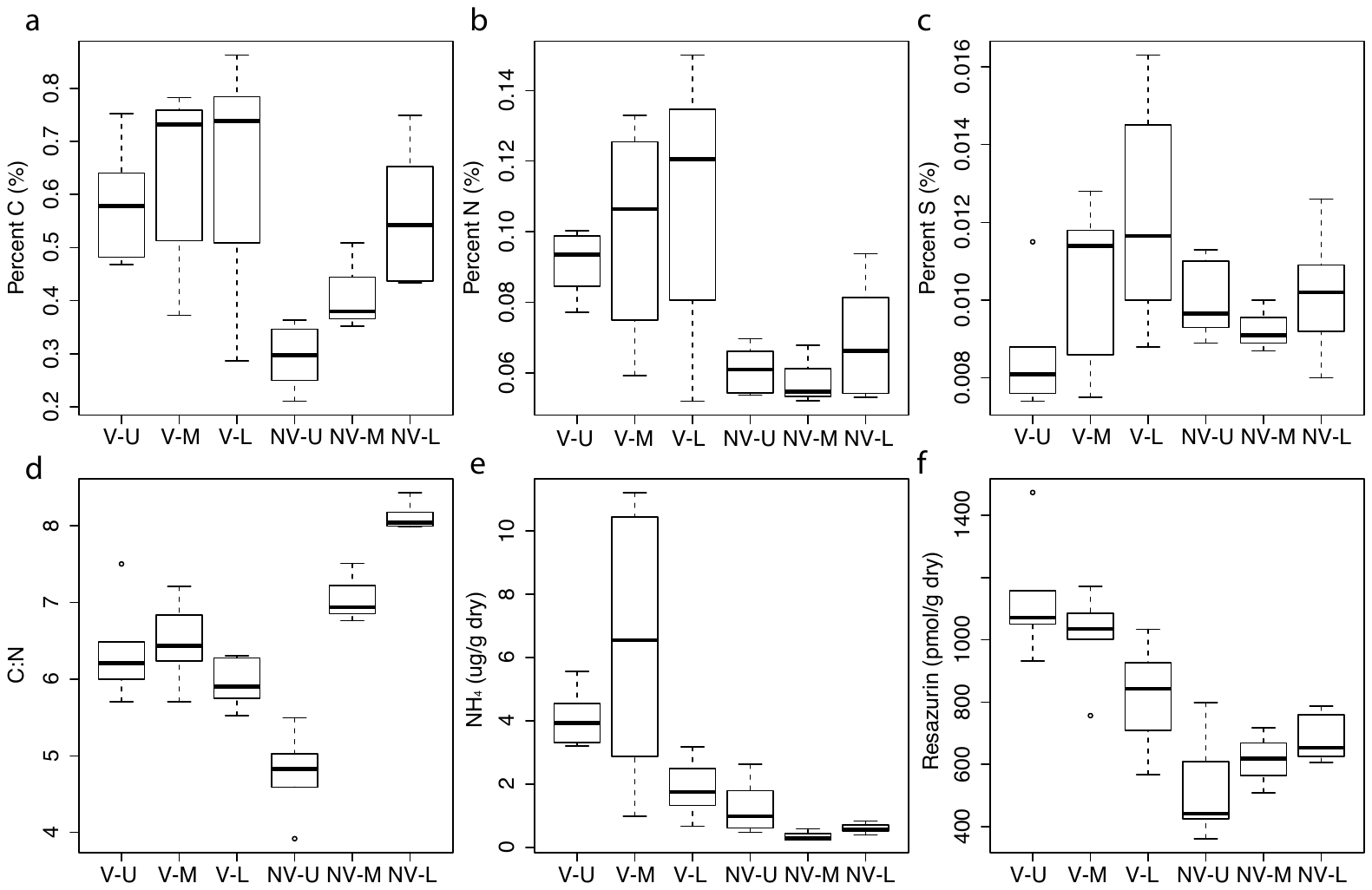

**Fig. S1. Boxplots of carbon, nitrogen, and aerobic metabolism in each sediment profile.** Each box represents all depths in a given core. ‘V’ and ‘NV’ denote a ‘vegetated’ or ‘unvegetated’ state. ‘U’, ‘M’, and ‘L’ represent upper, middle, and lower positions within the transect at each vegetation state. Upper and lower hinges of the box plots represent the 75^th^ and 25^th^ percentiles and whiskers represent 1.5 times the 75^th^ and 25^th^ percentiles, respectively. Five of seven hotspots are in V-U (depths 0-10, 20-30, 30-40, 40-50, and 50-60 cm), while the two remaining hotspots are in V-M (depths 20-30 and 30-40 cm). Low-activity sediments are distributed as follows: NV-U (depths 0-10, 30-40, and 40-50 cm); NV-M (depth 50-60 cm); NV-L (depths 0-10 and 20-30 cm); and V-L (depth 0-10 cm).

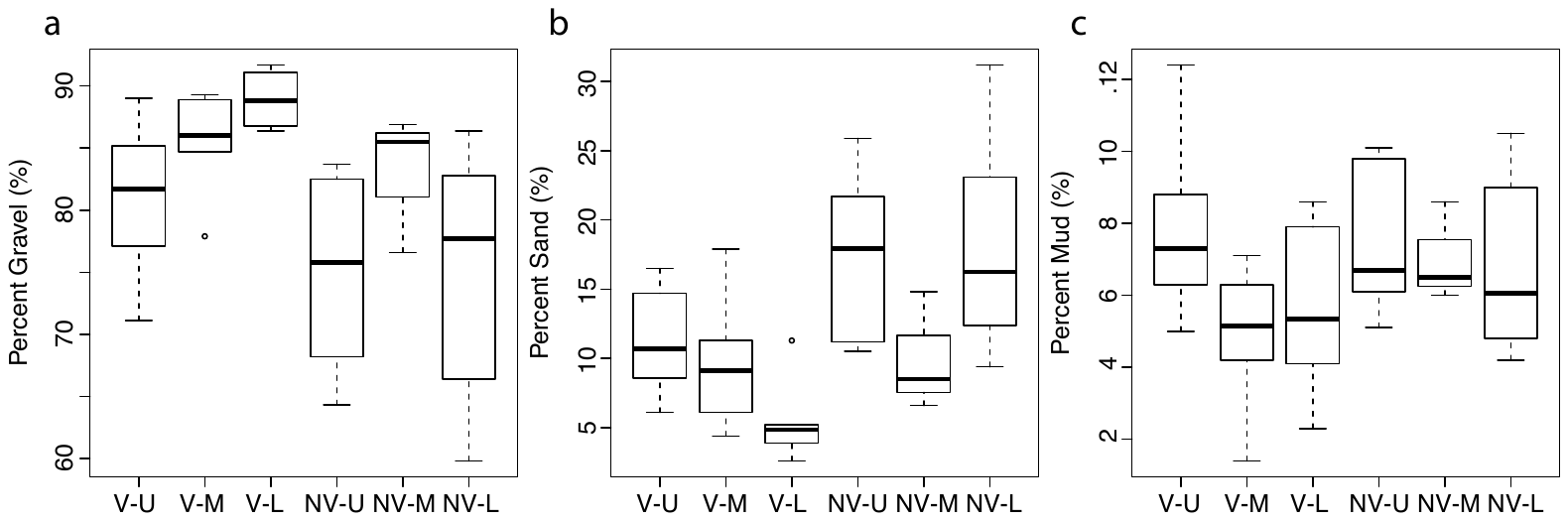

**Fig. S2. Boxplots of grain size distributions in each sediment profile.** Each box represents all depths in a given core. ‘V’ and ‘NV’ denote a ‘vegetated’ or ‘unvegetated’ state. ‘U’, ‘M’, and ‘L’ represent upper, middle, and lower positions within the transect at each vegetation state. Upper and lower hinges of the box plots represent the 75^th^ and 25^th^ percentiles and whiskers represent 1.5 times the 75^th^ and 25^th^ percentiles, respectively. Five of seven hotspots are in V-U (depths 0-10, 20-30, 30-40, 40-50, and 50-60 cm), while the two remaining hotspots are in V-M (depths 20-30 and 30-40 cm). Low-activity sediments are distributed as follows: NV-U (depths 0-10, 30-40, and 40-50 cm); NV-M (depth 50-60 cm); NV-L (depths 0-10 and 20-30 cm); and V-L (depth 0-10 cm).

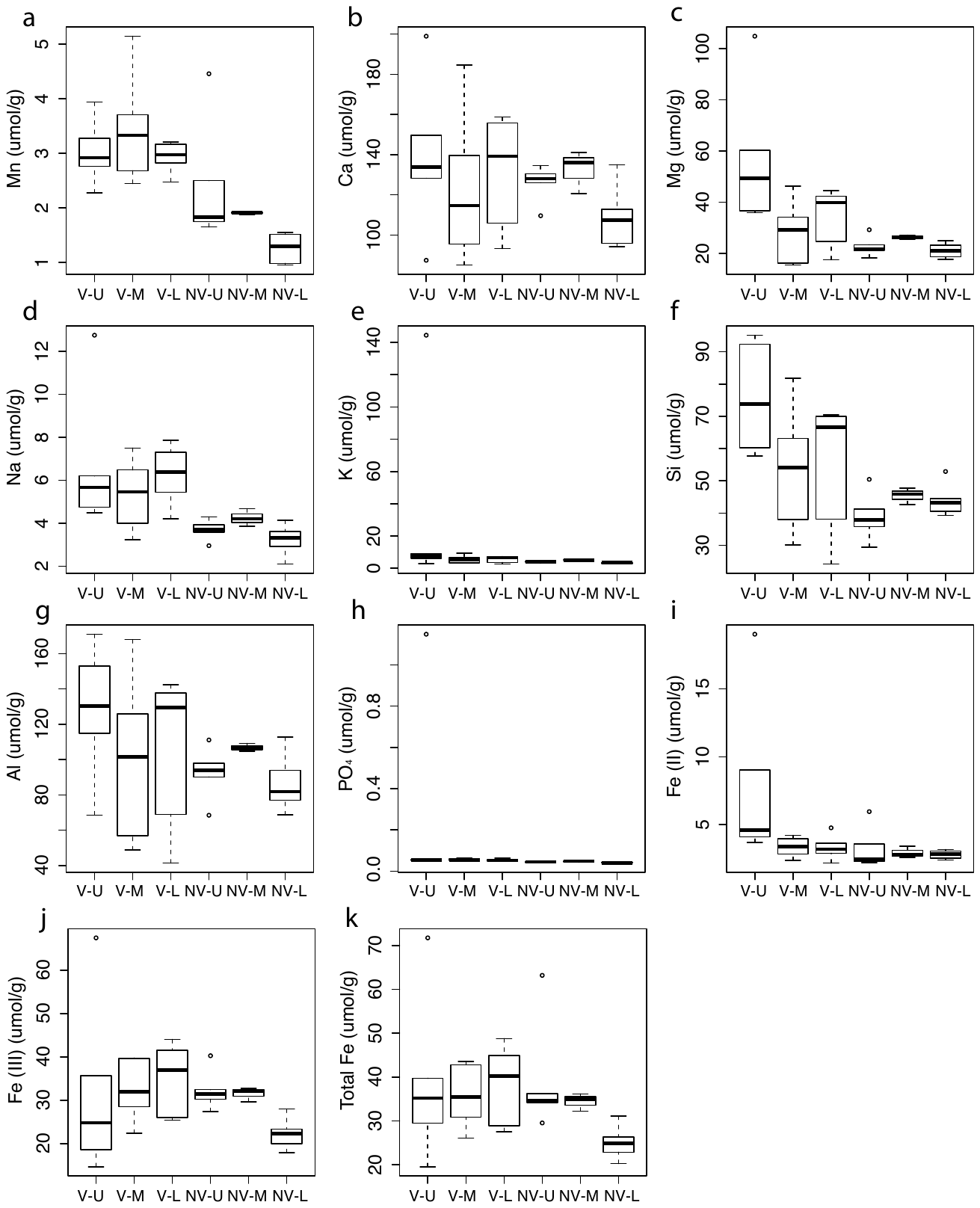

**Fig. S3. Boxplots of geochemistry in each sediment profile.** Each box represents all depths in a given core. ‘V’ and ‘NV’ denote vegetation state as in the main text. ‘U’, ‘M’, and ‘L’ represent upper, middle, and lower positions within the transect at each vegetation state. Upper and lower hinges of the box plots represent the 75^th^ and 25^th^ percentiles and whiskers represent 1.5 times the 75^th^ and 25^th^ percentiles, respectively. Five of seven hotspots are in V-U (depths 0-10, 20-30, 30-40, 40-50, and 50-60 cm), while the two remaining hotspots are in V-M (depths 20-30 and 30-40 cm). Low-activity sediments are distributed as follows: NV-U (depths 0-10, 30-40, and 40-50 cm); NV-M (depth 50-60 cm); NV-L (depths 0-10 and 20-30 cm); and V-L (depth 0-10 cm).

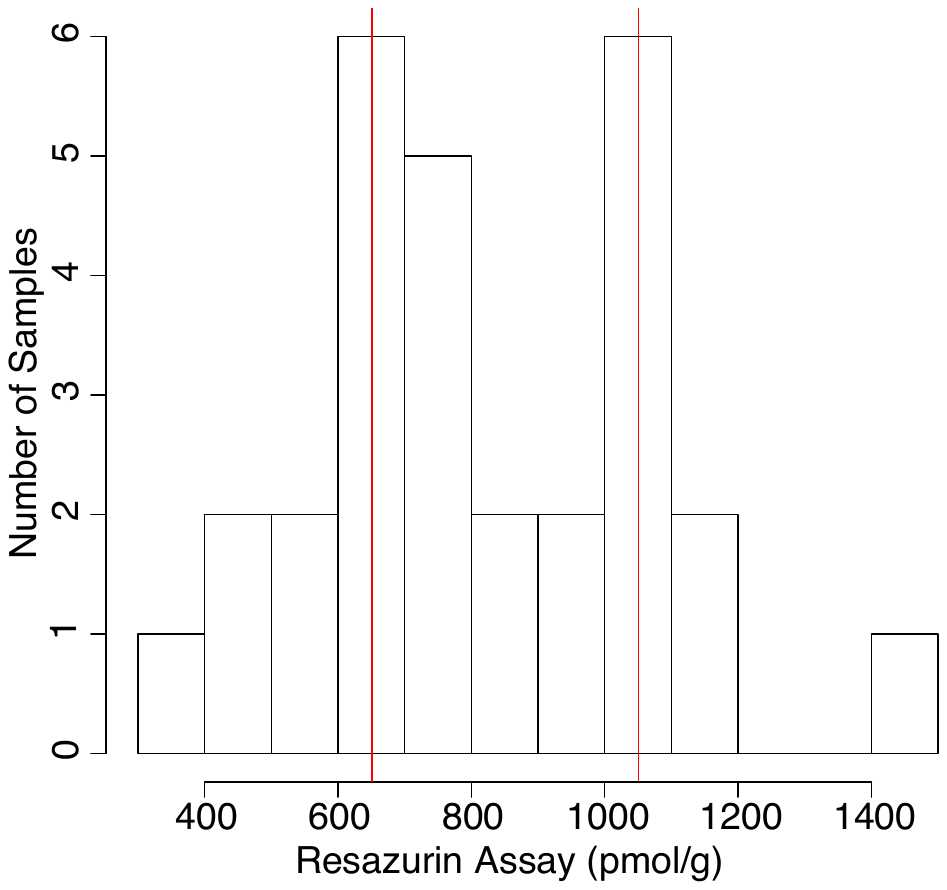

**Fig. S4. Histogram of Aerobic Metabolism Rates.** Red lines denote cut offs for low-activity (<619.45) and hotspot sediments (>1033.66).

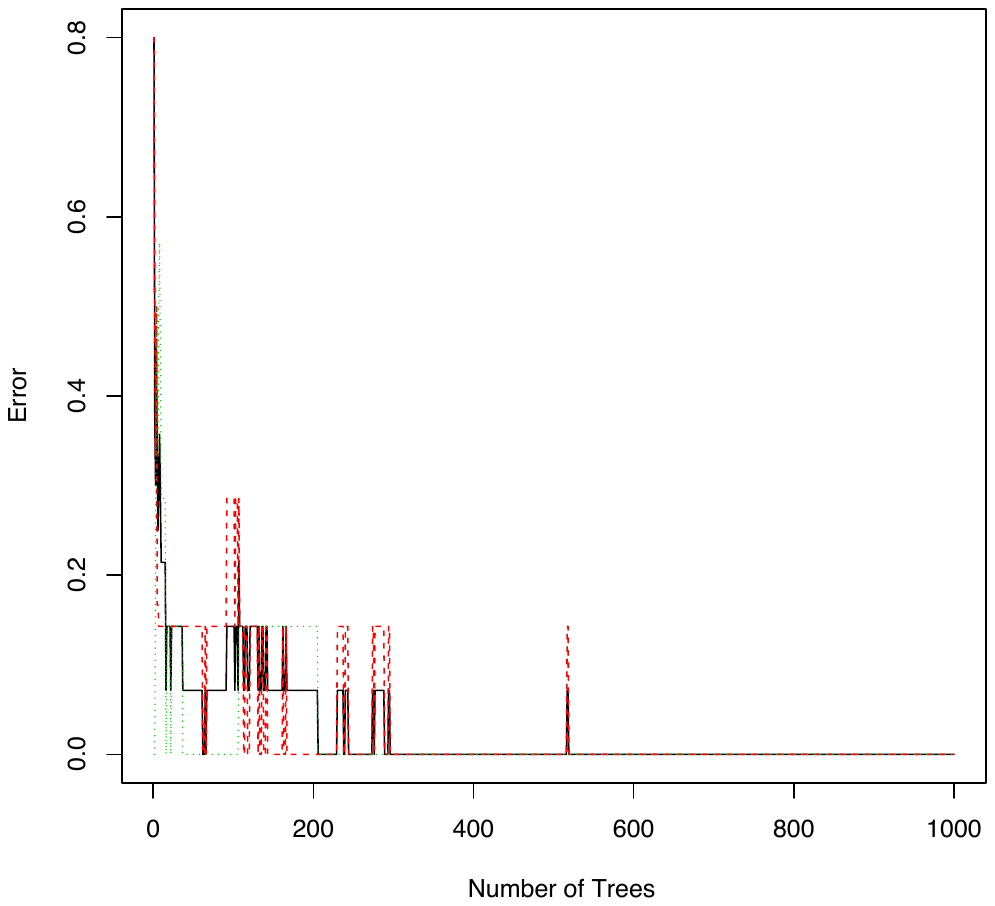

**Fig. S5. Random Forest Error Rate.** Error rates of random forests converged to near 0 over 1000 trees.

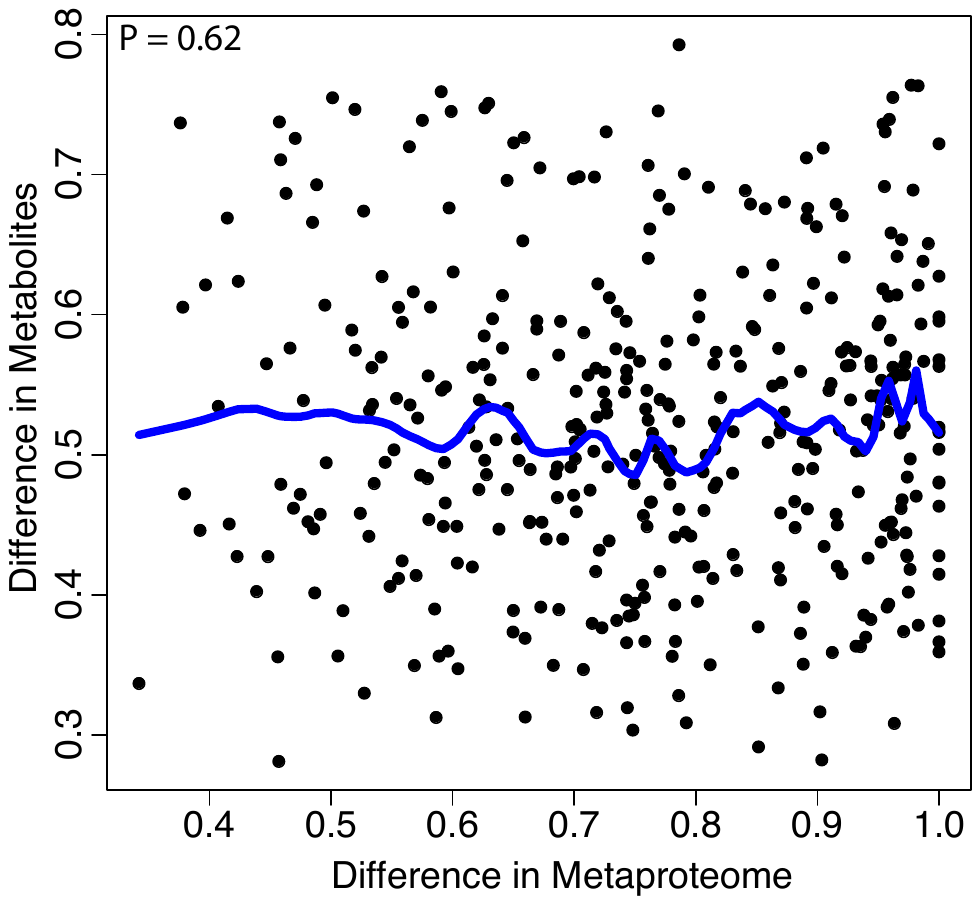

**Fig S6. Comparison of Metaproteome and Metabolome Composition.** Metabolite composition was determined using Sorenson dissimilarity, and metaproteome dissimilarity was determined with Bray-Curtis dissimilarity. The blue line depicts locally weighted scatterplot smoothing to visual data trends. The P-values is derived from a Mantel test of metabolome dissimilarity vs. metaproteome dissimilarity.

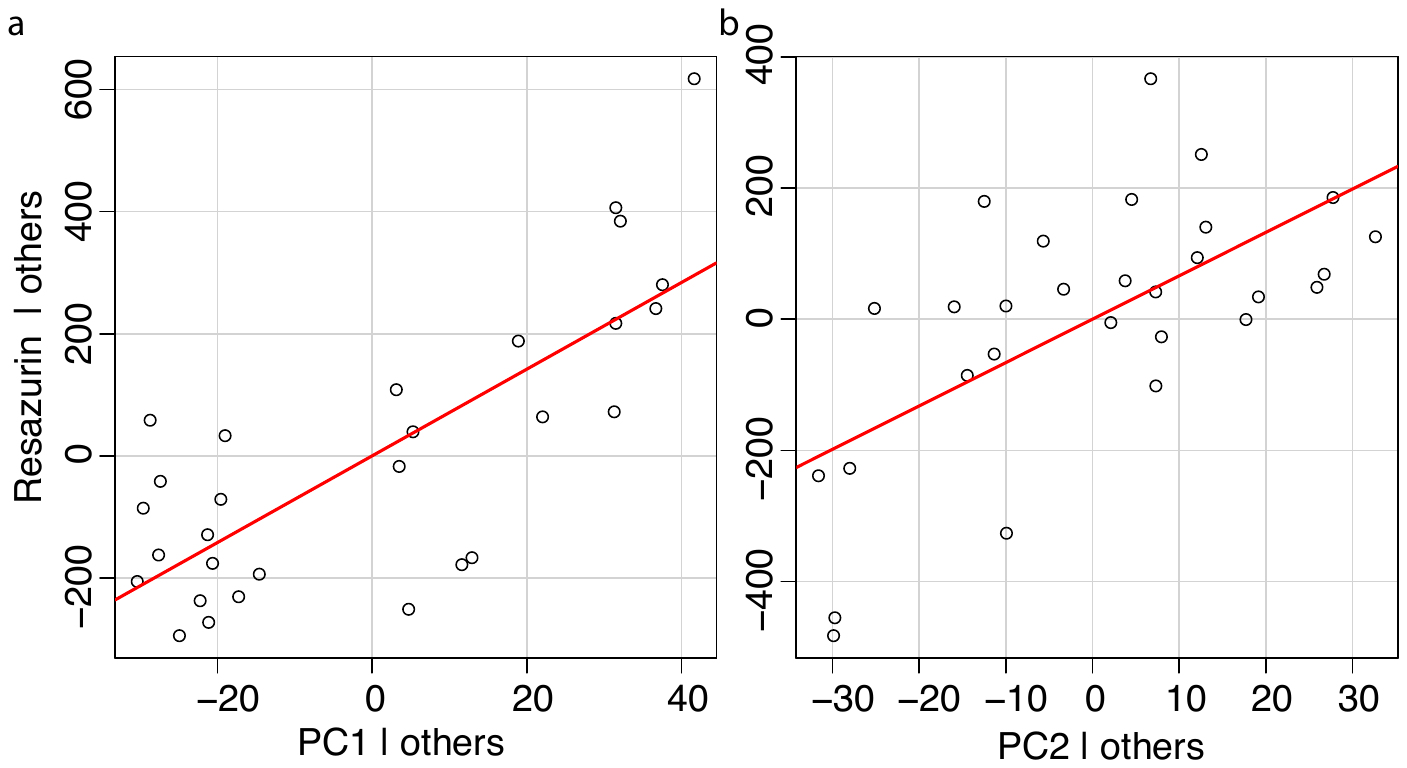

**Fig S7. Relationships of Metabolome PC1 and PC2 with Aerobic Metabolism.** (a) shows the correlation between PC1 and aerobic metabolism while controlling for PC2, and (b) shows the correlation between PC2 and aerobic metabolism while controlling for PC1. The contributions of each component are largely independent from each other.

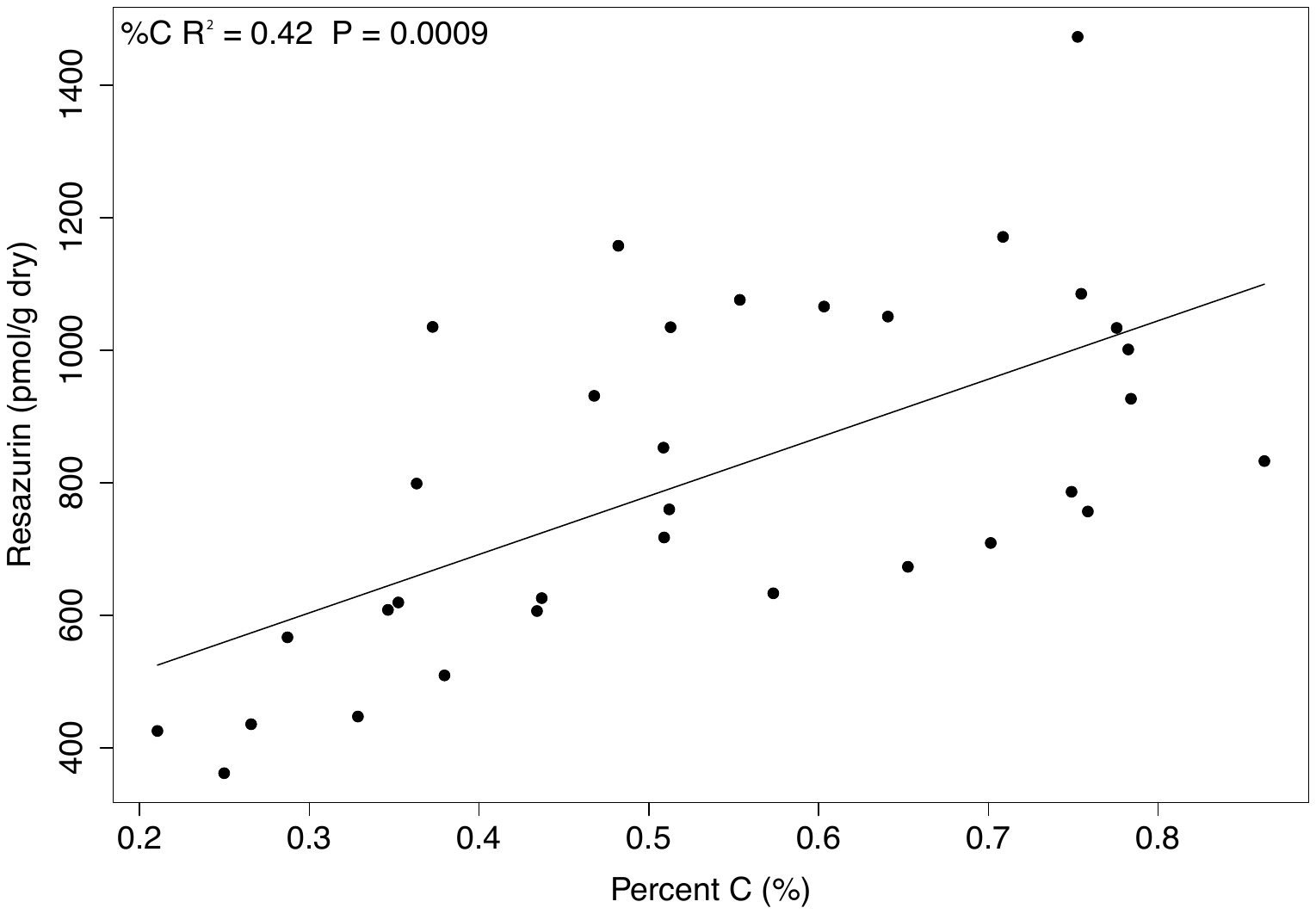

**Fig. S8. Relationship between Sediment C Content and Aerobic Metabolism.** Statistics are derived from a linear mixed effects model with %C as a fixed effect and depth as a random effect.

**Table S1. List of molecules used to identify biochemical transformations (i.e., masses gained or lost).**

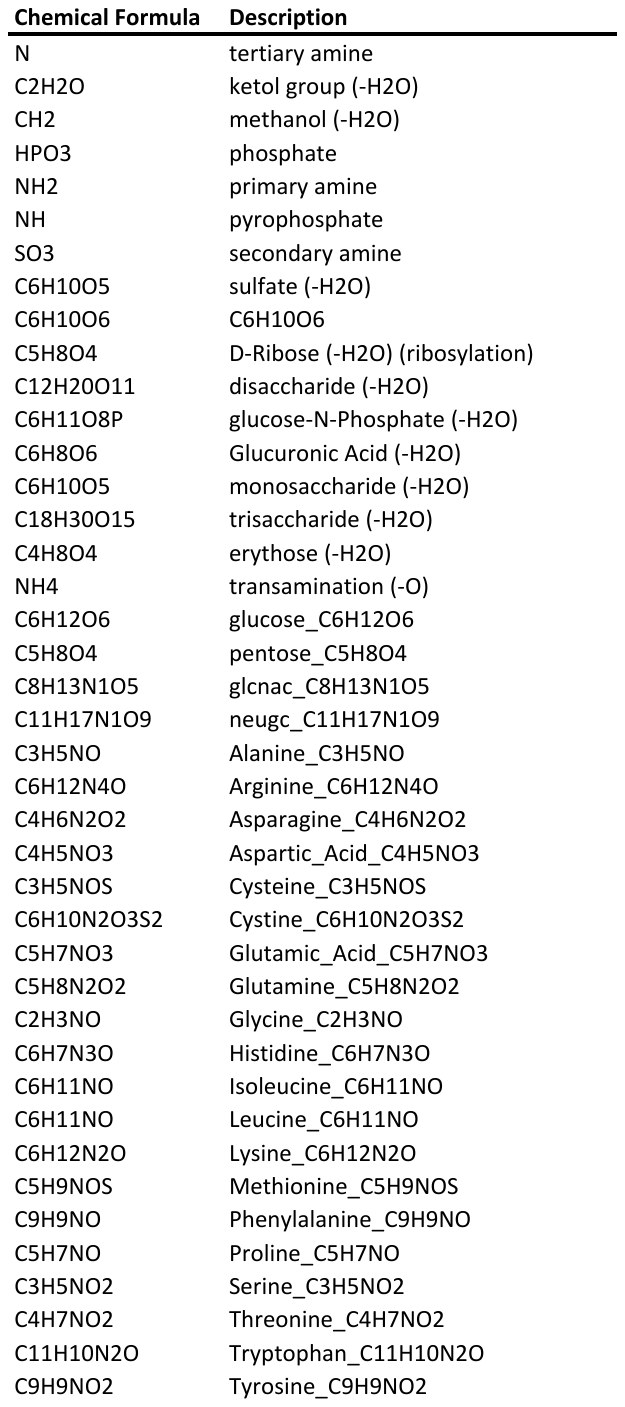

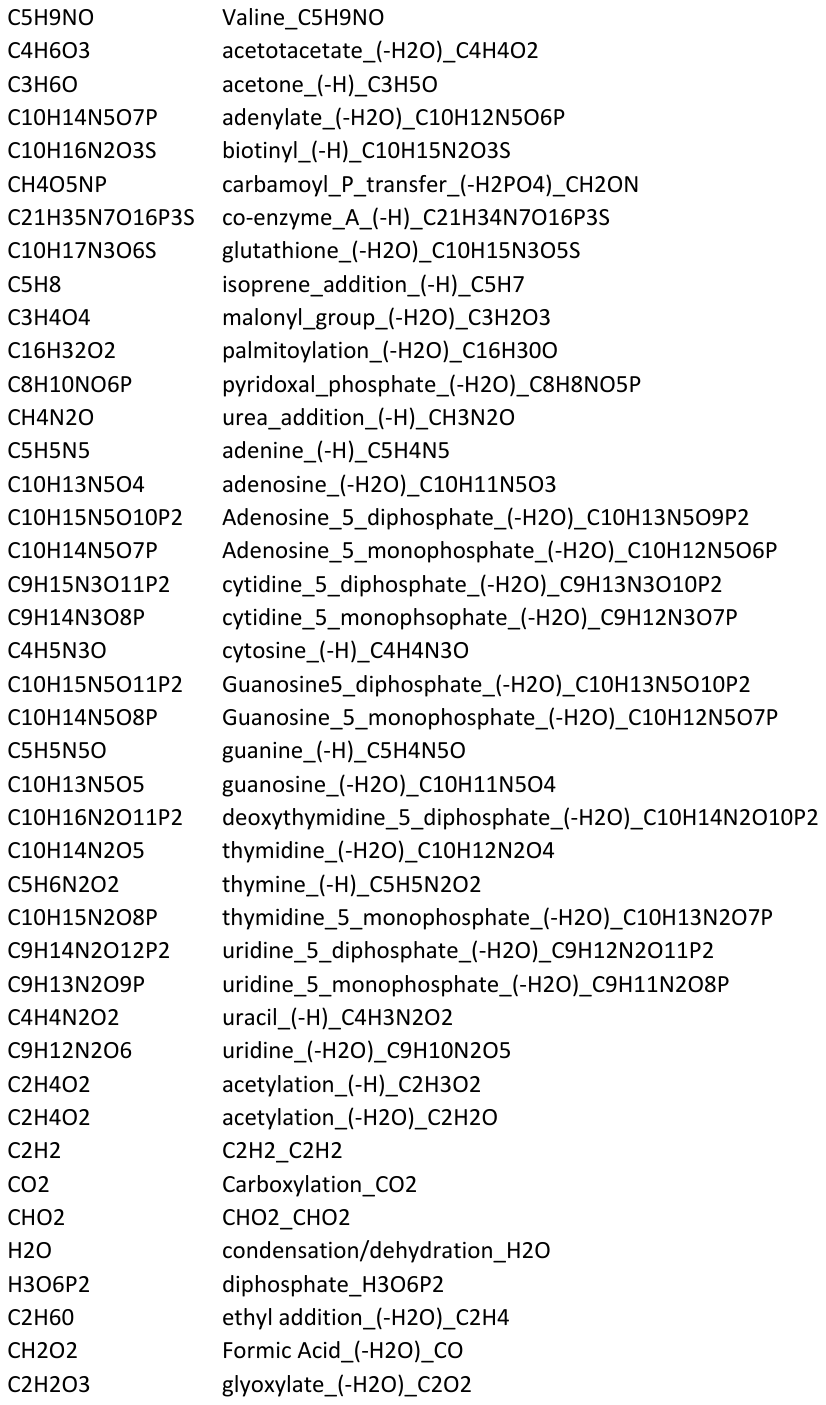

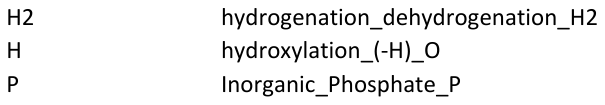

**Table S2. Information on Metagenomic Sequences and Assemblies.** Data are available at img.jgi.doe.gov.

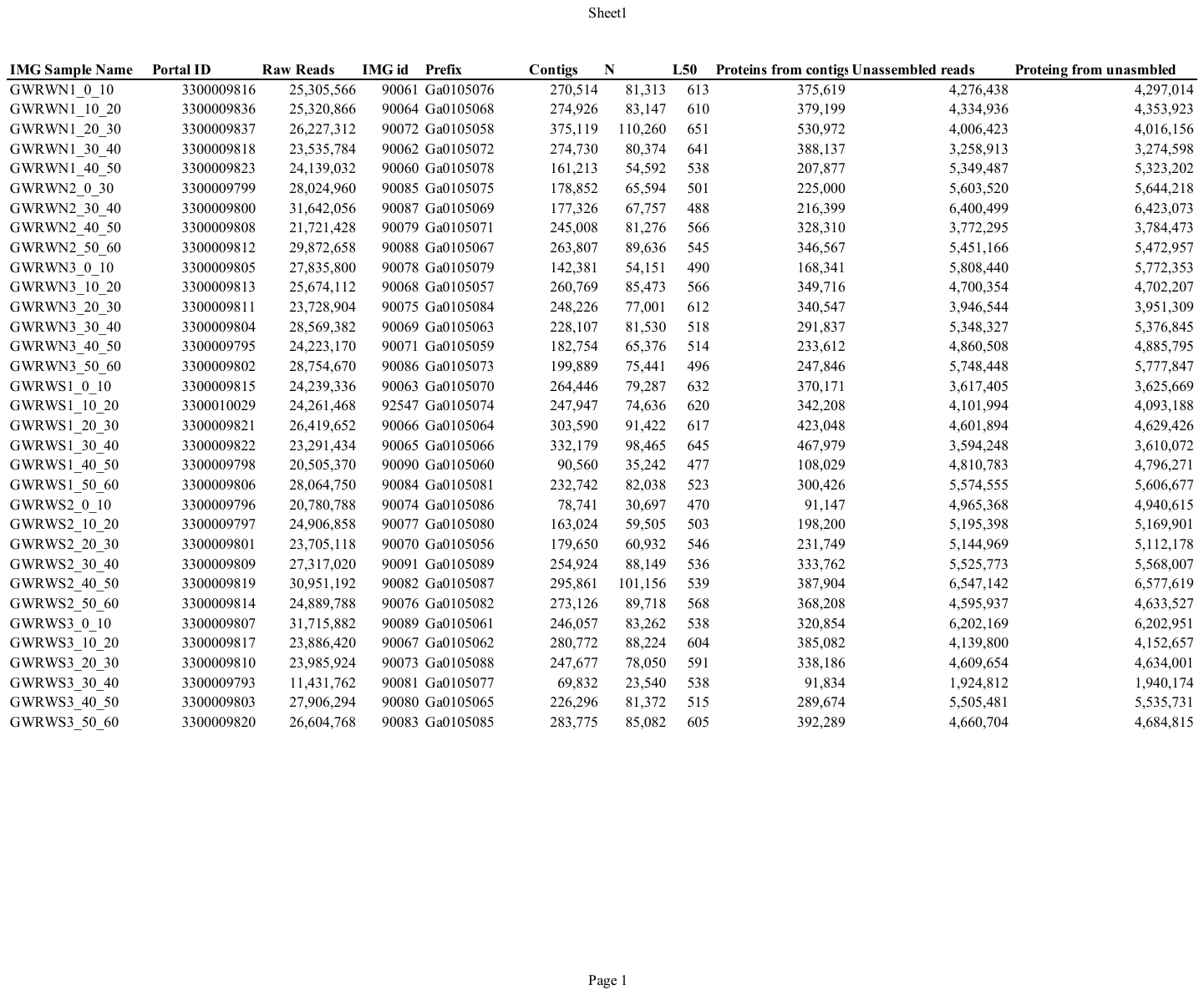

**Table S3. Nitrogenous transformations**

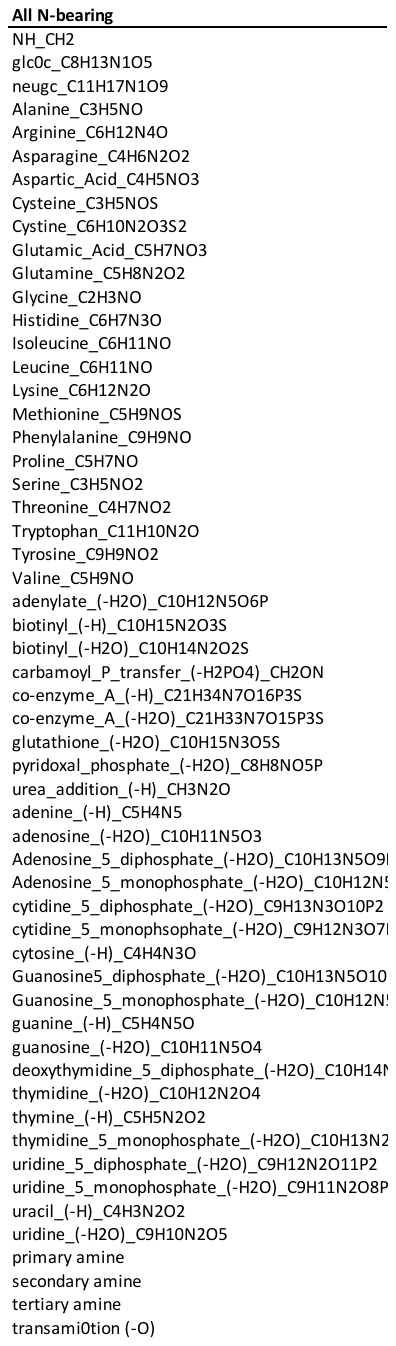

**Table S4. Transformations Not Involving N**

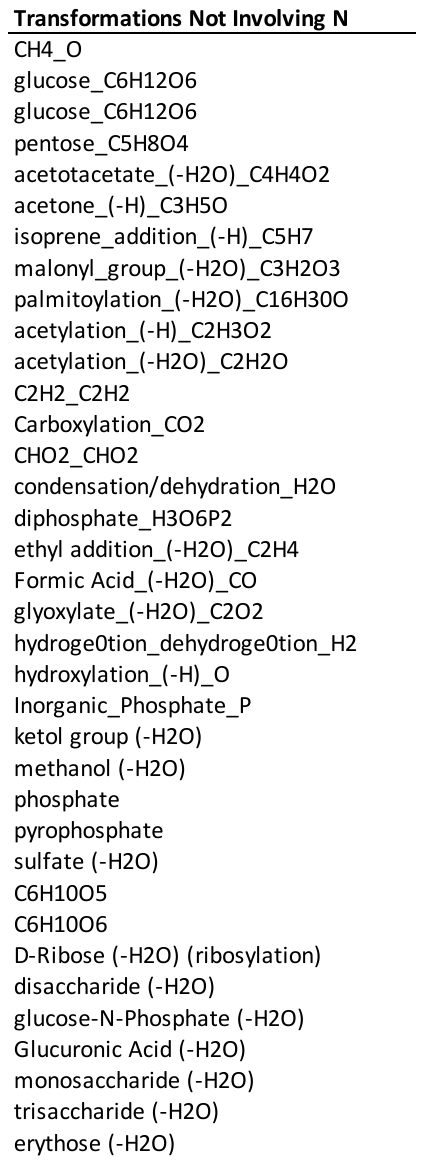

**Table S5. Amino Acid Transformations**

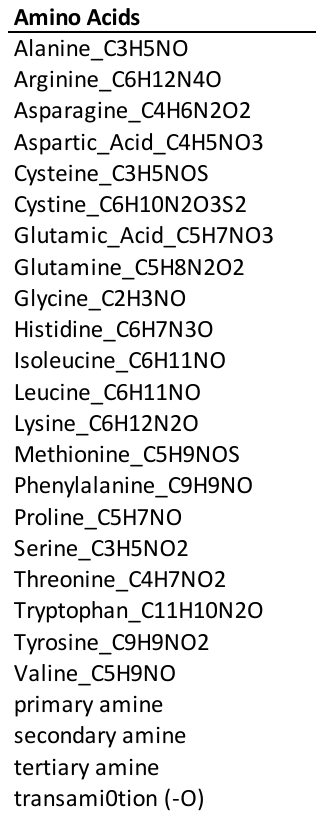

**Table S6. Complex N Transformations**

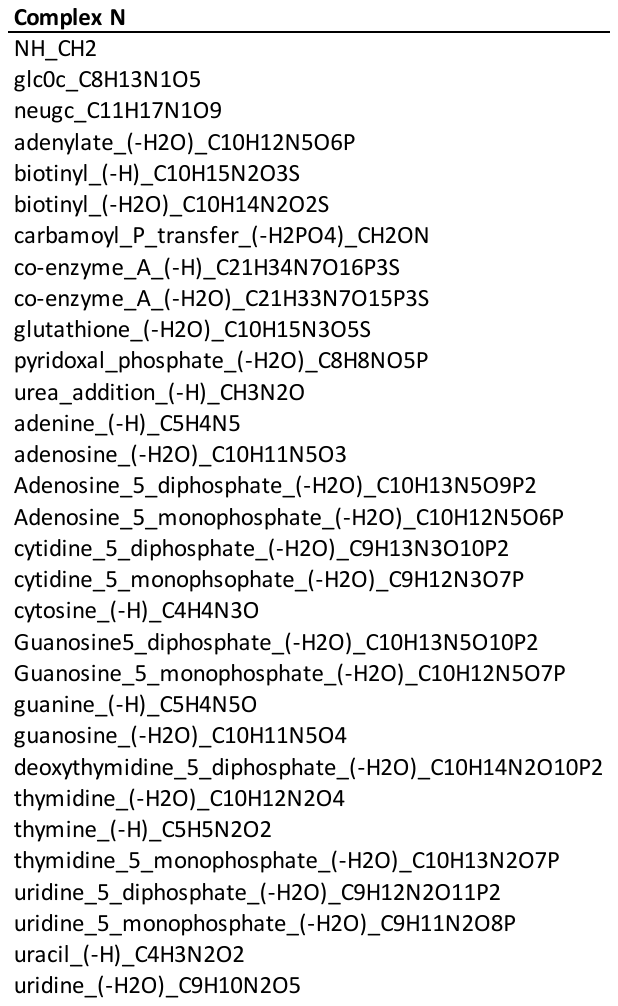

**References.**

Herzsprung P, Osterloh K, von Tümpling W, Harir M, Hertkorn N, Schmitt-Kopplin P, et al. Differences in DOM of rewetted and natural peatlands–Results from high-field FT-ICR-MS and bulk optical parameters. Science of The Total Environment 2017; 586: 770-781.

Kellerman AM, Kothawala DN, Dittmar T, Tranvik LJ. Persistence of dissolved organic matter in lakes related to its molecular characteristics. Nature Geoscience 2015; 8: 454-457.

Rossel PE, Bienhold C, Boetius A, Dittmar T. Dissolved organic matter in pore water of Arctic Ocean sediments: Environmental influence on molecular composition. Organic Geochemistry 2016; 97: 41-52.

Tfaily MM, Chu RK, Tolić N, Roscioli KM, Anderton CR, Paša-Tolić L, et al. Advanced solvent based methods for molecular characterization of soil organic matter by high-resolution mass spectrometry. Analytical chemistry 2015; 87: 5206-5215.

Tfaily MM, Chu RK, Toyoda J, Tolić N, Robinson EW, Paša-Tolić L, et al. Sequential extraction protocol for organic matter from soils and sediments using high resolution mass spectrometry. Analytica Chimica Acta 2017; 972: 54-61.

Ward CP, Cory RM. Chemical composition of dissolved organic matter draining permafrost soils. Geochimica et Cosmochimica Acta 2015; 167: 63-79.

Zhang L, Wang S, Xu Y, Shi Q, Zhao H, Jiang B, et al. Molecular characterization of lake sediment WEON by Fourier transform ion cyclotron resonance mass spectrometry and its environmental implications. Water Research 2016; 106: 196-203.
